# Transcriptome and chromatin accessibility mapping reveals a type I Interferon response triggered by *Mycobacterium tuberculosis* infection

**DOI:** 10.1101/2022.08.11.503537

**Authors:** Katrina Madden, Rayan El Hamra, Stefania Berton, Gonzalo G. Alvarez, Alexandre Blais, Jim Sun

**Author notes:** Correspondence to Jim Sun and Alexandre Blais.

## Abstract

Tuberculosis, a deadly infectious lung disease caused by *Mycobacterium tuberculosis* (Mtb), remains the leading cause of bacterial disease-related deaths worldwide. The success of Mtb as a human pathogen depends on its ability to manipulate host immune response pathways, many of which are regulated by epigenetic mechanisms that control the accessibility of chromatin to the transcriptional machinery. Recent reports suggest that host phosphatases, such as PPM1A, may play a role in the regulation of chromatin accessibility during bacterial infections. However, changes in genome-wide chromatin accessibility during Mtb infection and whether PPM1A plays a role in this process remains unknown. Using combinatorial chromatin accessibility (ATAC-seq) and transcriptomics (RNA-seq) profiling of wild-type (WT), PPM1A knockout (ΔPPM1A) and PPM1A overexpressing (PPM1A^+^) macrophages, we demonstrate that Mtb infection induces global chromatin remodeling consistent with changes in gene expression signatures. The strongest concordant chromatin accessibility and gene expression signature triggered by Mtb infection was enriched for genes involved in the type I interferon (IFN) signaling pathways. Modulation of PPM1A expression results in altered chromatin accessibility signatures during Mtb infection that are reflected in the total number, chromosome location and directionality of change. Transcription factor motif analysis revealed an enrichment for transcription factors involved in the type I IFN pathway during Mtb infection, including IRF4, MEF2A, and JDP2. In contrast, both deletion and overexpression of PPM1A produced unique transcription factor enrichment signatures linked to the genomic regions with altered chromatin accessibility. Our study demonstrates that altered type I IFN responses in Mtb-infected macrophages occurs as a result of genome-wide changes in chromatin accessibility, and that PPM1A likely plays a role in a subset of these signatures.

## INTRODUCTION

Tuberculosis (TB) is caused by *Mycobacterium tuberculosis* (Mtb) infection, and until the 2020 coronavirus pandemic, it was the leading cause of death from a single infectious agent, accounting for over 1.5 million deaths in 2021 (World Health Organization 2021). Mtb infects the innate immune cells of the lungs, and the success of infection depends on the re-programming of host cells to evade antibacterial responses, while simultaneously maintaining host cell viability (Roy *et al*. 2018; Looney *et al*. 2021). Host immune dysregulation in TB has been demonstrated at the gene expression, or epigenetic level, which is frequently governed by the accessibility of chromatin to the transcriptional machinery (3–6). The most common epigenetic mechanisms that regulate gene expression through modulating accessibility of DNA to the transcriptional machinery include DNA methylation, conformation of nucleosomes and histones, and transcription factor (TF) binding. Genome-wide analyses of the DNA methylome of Mtb-infected cells compared to non-infected cells have demonstrated that changes in gene expression are mediated in part by gain or loss of DNA methylation (Zheng *et al*. 2016; Chen *et al*. 2020; Karlsson *et al*. 2021; Looney *et al*. 2021). Genome-wide histone methylation and acetylation patterns, such as H3K27ac, H3K4me3, H4K20me, H3K14ac or H3K27me2/3, have also been analyzed using ChIP-seq in macrophages infected with Mtb, *Mycobacterium bovis*, or blood cells from TB patients (Holla *et al*. 2016; Chen *et al*. 2017; Singh *et al*. 2017; Hall *et al*. 2019; Subuddhi *et al*. 2020; Del Rosario *et al*. 2022). These studies revealed that Mtb infection or TB disease induced many changes to nucleosome structure that altered gene expression to dampen antibacterial responses. The limitation of these studies is that ChIP-seq analyzes specific changes to nucleosome structure and does not simultaneously capture genome-wide changes to chromatin accessibility. The only previous report to investigate global chromatin accessibility patterns in macrophages upon Mtb infection was a report by Correa-Macedo et al. conducted in the context of HIV co-infection, which did not focus in-depth on the effects of Mtb infection alone (Correa-Macedo *et al*. 2021). In contrast, numerous studies have examined the impact of Mtb infection on the transcriptome of macrophages (Madden *et al*. 2022). These studies have shown a host-detrimental type I interferon (IFN) signature in Mtb infected cells and TB patients (Moreira-Teixeira *et al*. 2018).

Several reports have also documented the dysregulated expression of enzymes that mediate histone modifications in macrophages after Mtb infection, such as histone deacetylases HDAC1, HDAC3, and Sirtuins (SIRT) SIRT1, SIRT2, and SIRT3 (Chandran *et al*. 2015; Cheng *et al*. 2017; Moores *et al*. 2017; Kim *et al*. 2019; Bhaskar *et al*. 2020; Campo *et al*. 2021; Madhavan *et al*. 2021; Smulan *et al*. 2021). Changes to nucleosome structure that occur during pathogen invasion usually correlated to impaired antibacterial responses, and could be rescued by therapeutically targeting these enzymes (Chandran *et al*. 2015; Cheng *et al*. 2017; Moores *et al*. 2017; Kim *et al*. 2019; Bhaskar *et al*. 2020; Campo *et al*. 2021; Madhavan *et al*. 2021; Smulan *et al*. 2021). Similar mechanisms are employed by other intracellular pathogens, such as *Listeria monocytogenes*. Successful *L. monocytogenes* infection requires activation of SIRT2 by the host Protein Phosphatase Mg^2+^/Mn^2+^-dependent 1A (PPM1A) to deacetylate H3K18, which leads to disabled antibacterial immune responses (Pereira *et al*. 2018: 2). Interestingly, we have also reported that PPM1A is upregulated in macrophages during Mtb infection to disable multiple antibacterial response pathways including apoptosis, autophagy and inflammation (Sun *et al*. 2016; Schaaf *et al*. 2017; Smith *et al*. 2018; Berton *et al*. 2022a). Whether the host-detrimental effects of PPM1A activity during Mtb infection extend to the modulation of nucleosome structure or chromatin accessibility – via histone modifying enzymes or other mechanisms – is unknown.

Here, we use ATAC-seq and RNA-seq to determine the genome-wide changes that occur at the epigenetic (chromatin accessibility) and transcriptomic level in Mtb-infected human macrophages. We identify specific changes to chromatin accessibility and gene expression that are mediated by PPM1A expression. Extensive bioinformatic analysis revealed distinct chromatin accessibility signatures in Mtb-infected wild-type (WT) and PPM1A knockout (ΔPPM1A) macrophages compared to noninfected controls, and that these changes to chromatin accessibility were exacerbated in PPM1A overexpressing (PPM1A^+^) macrophages. The differential chromatin accessibility and gene expression patterns were highly enriched in genes associated to the type I IFN signaling pathways. A subset of type I IFN response genes where chromatin accessibility changed displayed striking and concordant changes to gene expression. Furthermore, genomic regions where accessibility was altered by Mtb infection were enriched for the binding sites of transcription factors involved in type I IFN responses and TB pathogenicity.

## RESULTS

### Generation of ATAC-seq libraries from Mtb-infected macrophages

To investigate the impact of Mtb infection on genome-wide chromatin accessibility patterns and subsequent transcriptional changes, we performed parallel ATAC-seq and RNA-seq using wild-type (WT) and genetically modified human THP-1 macrophages infected with Mtb mc^2^6206 (**Fig. 1A**). THP-1 macrophages are routinely used to study innate immune responses to Mtb infection because they are genetically tractable and reproduce key physiological features of primary human monocyte derived macrophages (hMDM) (Tsuchiya *et al*. 1980; Daigneault *et al*. 2010; Bosshart and Heinzelmann 2016; Forrester *et al*. 2018). Mtb mc^2^6206 is a derivative of virulent Mtb H37Rv (Jain *et al*. 2014) that retains all virulence genes and has been shown to elicit a comparable host immune response and susceptibility to TB antibiotics, and intra-macrophage replication and survival compared to its parent strain (Schaaf *et al*. 2017; Mouton *et al*. 2019). We determined that Mtb infection for 48 h at a multiplicity of infection (MOI) of 10 achieved the optimal infection conditions as determined by the highest percentage of infection (70%) and lowest cell death (10%), which can interfere with gene regulation **(Fig. S1)**. To determine chromatin accessibility patterns, we used the assay for transposase-accessible chromatin sequencing (ATAC-seq), which uses a hyperactive Tn5 transposase to mark by tagmentation, DNA that is relatively accessible (Corces *et al*. 2017). Biological duplicates of Mtb-infected and non-infected THP-1 macrophages (WT, ΔPPM1A, PPM1A^+^) were processed for ATAC-seq library generation. The OMNI-ATAC version of the ATAC-seq protocol was used to minimize mitochondrial genome representation in DNA libraries (Corces *et al*. 2017). We maximized the quality of sequencing reads by incorporating dual-end barcoding into the library design, which improves traditional single-end barcoding by limiting barcode hopping and increasing multiplexing (Kircher, Sawyer and Meyer 2012). Quality control of our ATAC-seq libraries demonstrated enrichment of loci known to be accessible or inaccessible in THP-1 cells at baseline **(Fig. S2A)**. Prior to sequencing, the quality of the pooled sample library was determined through bioanalyzer analysis where the size of DNA fragments on average were 52 bp and the concentration was 1.57 ng/μl **(Fig. S2B)**. Following sequencing, quality control analysis confirmed that average Tn5 transposome insert sizes ranged from 100-1000 bp **(Fig. S2C)** and that peaks were enriched in regions 5000 bp up or downstream of the transcriptional start site (TSS) **(Fig. S2D)**. We applied false discovery rate (FDR) cutoffs of [FDR] < 0.05, and peaks and were retained if they were present in at least two samples. The number of consensus peaks found in at least two samples was 317,537 peaks per sample. Principal component analysis (PCA) of read coverage over all peaks demonstrated that genotype and infection status were the largest and second largest contributor to variance in the dataset, respectively (**Fig. S2E)**. Peaks from biological replicates also clustered together, confirming the reproducibility of our data.

**Figure 1.**
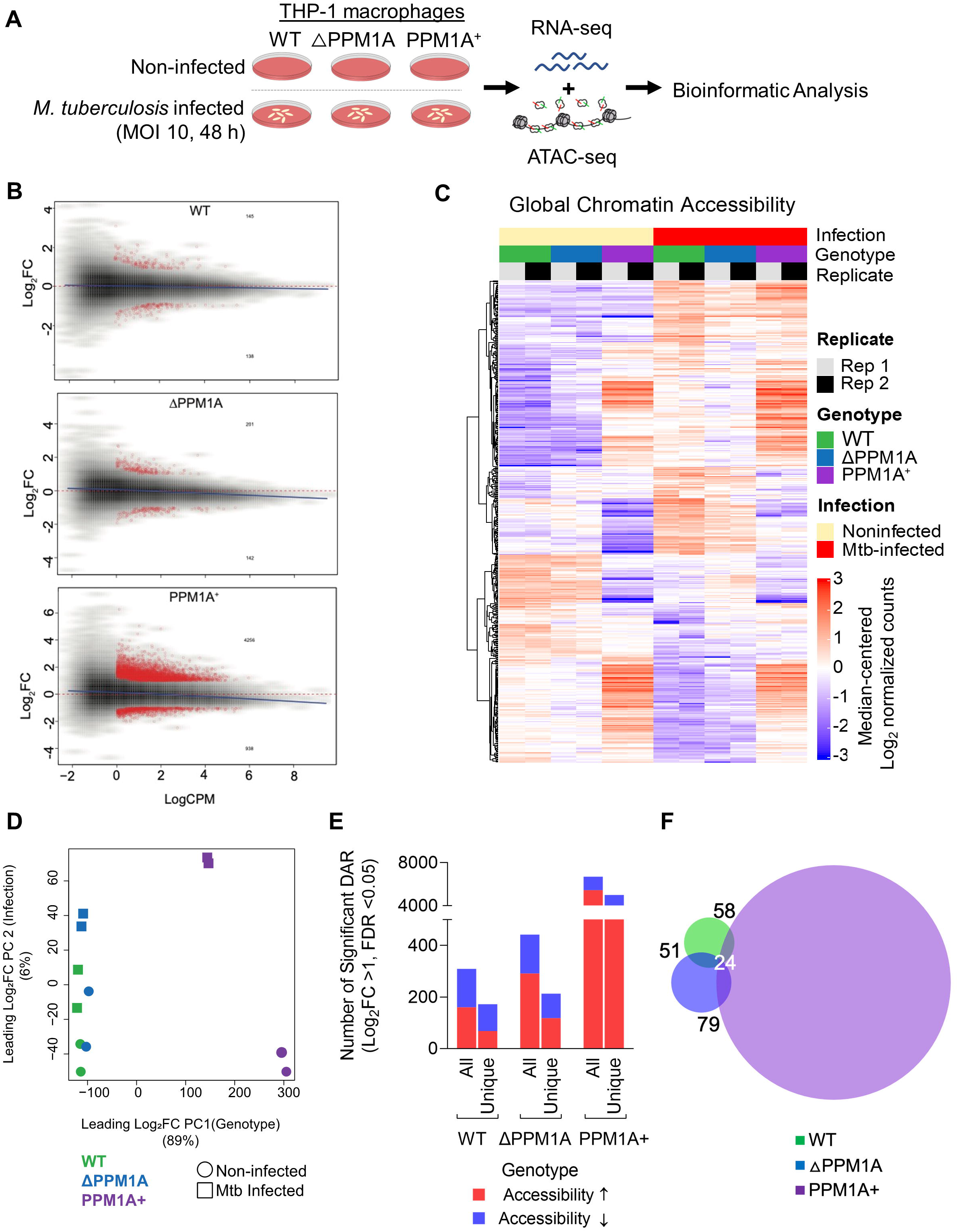
Mtb infection induces global chromatin remodeling in macrophages. **(A)** Experimental schematic. WT, ΔPPM1A, and PPM1A^+^ THP-1 macrophages were infected with Mtb mc^2^6206 at an MOI of 10 for 48 h. Then, Mtb-infected and non-infected control macrophages were harvested for ATAC-seq (biological duplicates) and RNA-seq (biological triplicates) in parallel from the same wells. **(B)** MA plots showing log_2_ fold-change versus average signal (log of counts per million, LogCPM) in Mtb-infected samples compared to controls, in each of three genotypes studied. Peaks with significant chromatin accessibility (CA) change are marked in red. Blue lines represent the general signal trend by loess fit. Red dotted lines mark the location of identical signal. **(C)** Heatmap of CA in regions that were significantly differentially accessible in WT cells upon Mtb infection. CA in WT, ΔPPM1A and PPM1A^+^ macrophages before and after Mtb infection is shown. All ATAC-seq peaks were retained if they had greater than 24 counts in at least two samples. Differential peaks between Mtb-infected WT macrophages and non-infected controls with absolute value of log_2_FC > 1 and FDR < 0.05 were considered significant. **(D)** Principal component analysis (PCA) showing significant differential ATAC-peaks separate and cluster together based on the first and second principal components which are genotype and Mtb infection status, respectively. Genotype accounts for 89% of the variance observed in all significant DAR, and Mtb infection accounts for 6%. **(E)** Total and unique DAR. The first bar displays the total number of all significant differential peaks in each genotype after Mtb infection, separated by whether CA increased or decreased. The second bar displays the total number of peaks that are unique to each genotype, separated by whether CA increased or decreased. **(F)** Venn diagram showing the overlap of significant differential peaks between Mtb-infected WT, ΔPPM1A and PPM1A^+^ macrophages and non-infected macrophages.

### Mtb infection induces global chromatin remodeling in macrophages

Differential analysis of DNA accessibility at peaks showed significant genome-wide changes to chromatin accessibility in WT, ΔPPM1A and PPM1A^+^ cells after Mtb infection **(Fig. 1B)**. Chromatin accessibility profiles in ΔPPM1A and PPM1A^+^ cells diverged from WT cells in many regions, before and after Mtb infection **(Fig. 1C)**. There was also a marked increase in the number of significant differentially accessible regions (DAR) in PPM1A^+^ cells compared to both WT and ΔPPM1A cells **(Fig. 1B, C)**. PCA of peaks that are differentially accessible in any of the pairwise comparisons demonstrated that all differential peaks also clustered based on genotype and infection **(Fig. 1D)**, resembling the PCA of all peaks (**Fig. S2E)**. Genotype accounted for 89% of the variance observed, while infection accounted for 6%. In WT macrophages, Mtb infection induced 309 significant DAR, where accessibility increased in 160 regions (52%) and decreased in 149 regions **(Fig. 1E)**. Of these loci, 266 (86%) annotated to genes based on their distance from the TSS. The number of significant DAR upon Mtb infection in ΔPPM1A cells increased to 442, where accessibility increased in 291 regions (66%) and decreased in 151 regions **(Fig. 1E)**. In ΔPPM1A cells, the proportion of regions where accessibility increased was 14% greater than in WT cells, and 364 (82%) peaks annotated to genes in ΔPPM1A cells. The number of significant DAR after Mtb infection substantially increased in PPM1A^+^ cells to 6702 loci, which represents a 22-fold increase in total significant DAR compared to WT cells. The proportion of regions where accessibility increased was higher in PPM1A^+^ cells, where accessibility increased in 5443 regions, representing 81% of total DAR **(Fig. 1E)**. The accessibility decreased in 1259 regions in PPM1A^+^ cells and 5372 (80%) of peaks annotated to genes. While all three genotypes shared a set of conserved DAR induced by Mtb infection, each genotype also had a set of unique DAR. There were 172 unique peaks in WT macrophages, where accessibility increased in 68 and decreased in 104 regions. In ΔPPM1A cells, there were 213 total unique peaks, where accessibility increased in 118 and decreased in 95 regions **(Fig. 1E)**. In PPM1A^+^ cells, there were 4994 unique peaks where accessibility increased in 3999 and decreased in 995 peaks **(Fig. 1E)**. 57 and 51 DAR were shared between WT and ΔPPM1A or PPM1A^+^ cells respectively, and 79 were shared between ΔPPM1A and PPM1A^+^ cells **(Fig. 1F)**. There were 24 DAR shared between all three genotypes. In WT, ΔPPM1A, and PPM1A^+^ cells, 16%, 14% and 13% of significant DAR annotated to promoter regions, while the majority (32-35%) of significant DAR in all genotypes annotated to regions labelled as ‘other introns’ **(Fig. S2F)**.

### Mtb induced chromatin accessibility signatures are enriched in type I IFN signaling pathways

To identify biological pathways and processes that are associated with Mtb-infection induced changes to chromatin accessibility, we performed GO analysis of DAR using the genomic regions enrichment annotations tool (GREAT) (McLean *et al*. 2010). All significant DAR in WT, ΔPPM1A and PPM1A^+^ macrophages upon Mtb infection were analysed separately with GREAT. DAR were annotated to genes based on the association rules provided by GREAT (**Fig. S3A, B, C**). In each case, GO terms with hypergeometric adjusted p-values < 0.05 were ranked by enrichment and top-scoring non-redundant terms with at least 10 annotates genes were plotted (**Fig. 2A**). The GO terms reported in Figure 2A combine the most highly enriched GO terms associated to DAR upon Mtb infection in WT, ΔPPM1A, and PPM1A^+^ cells. The most highly enriched GO terms associated to DAR in all genotypes were dominated by type I interferon responses (**Fig. 2A**). However, type I IFN response pathways were significantly more enriched in DAR in WT and ΔPPM1A cells, than in DAR in PPM1A^+^ cells. Mtb-infected WT cells showed four unique enriched GO terms, whereas there was only one unique enriched GO term in ΔPPM1A cells, and none in PPM1A^+^ cells (**Fig. 2B**). GO terms that were unique to WT cells included “2’ deoxyribonucleotide biosynthetic process” and related terms that were enriched by DAR annotated to *AK5* and *CMPK2*. In ΔPPM1A cells, “Positive regulation of neutrophil chemotaxis” was the sole unique GO term and was enriched by DAR annotated to *IL-23, CXCL2, CXCL5* and *CXCL8* (**Fig. 2B).**

**Figure 2.**
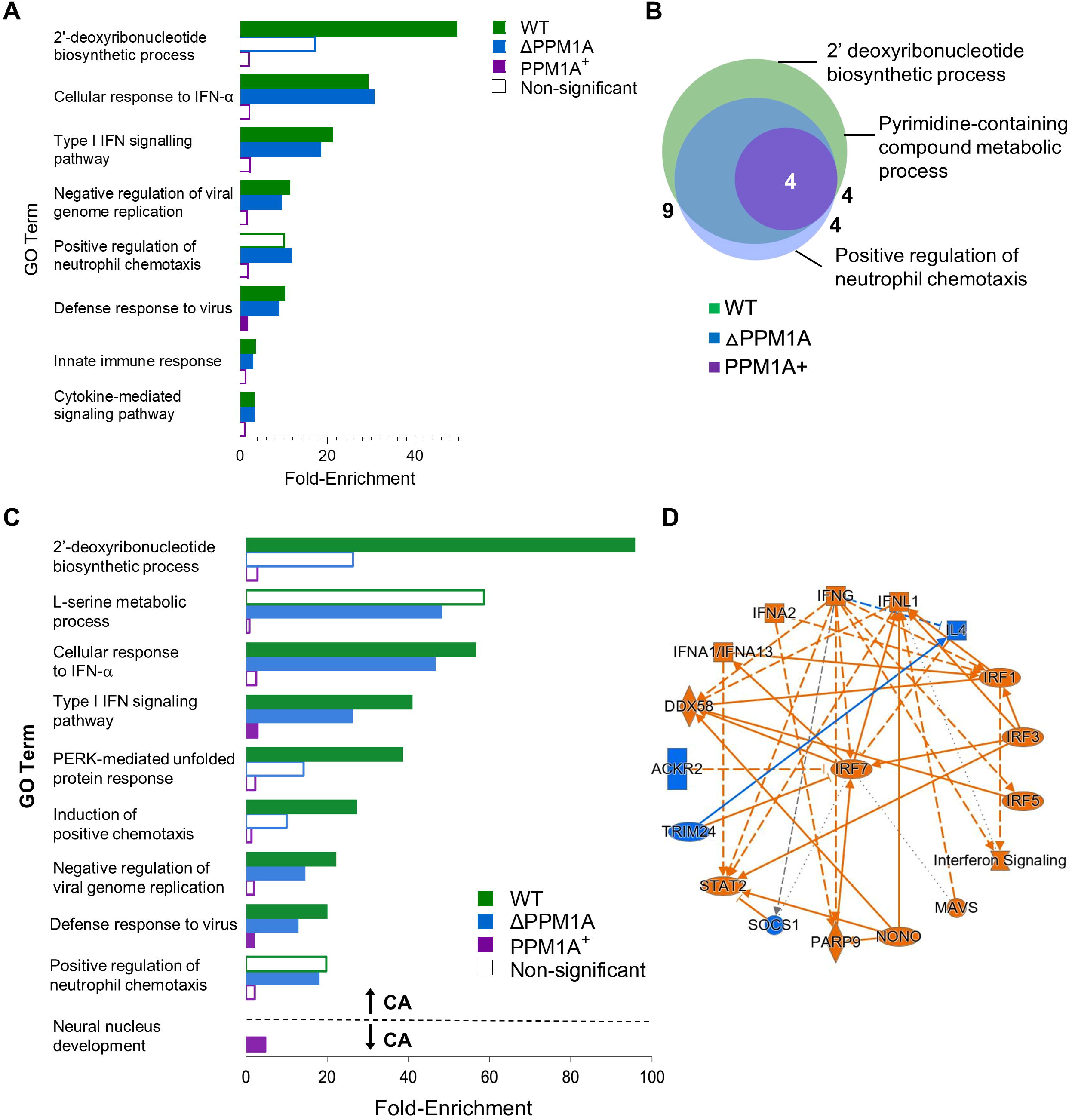
Mtb infection induces changes to chromatin accessibility that are enriched in type I IFN regulatory pathways. **(A)** Gene ontology (GO) term enrichment analysis was conducted using GREAT with significant differential peaks in WT, ΔPPM1A and PPM1A^+^ macrophages following Mtb infection. GO terms with FDR < 0.05 and at least 10 annotated genes were considered significant. GO terms were ranked by enrichment and the top 5 non-redundant terms from each genotype were included. Many of the top 5 non-redundant terms overlapped between genotypes. (**B)** Venn diagram displaying common significant GO terms between WT, ΔPPM1A and PPM1A^+^ macrophages after Mtb infection. **(C)** Separate GO enrichment analysis was conducted for chromatin regions where accessibility increased or decreased. Significant DAR in each genotype were split into two groups based on whether CA increased (log_2_FC > 1) or decreased (log_2_FC < 1). GO terms were ranked and plotted as in (A). The top 5 non-redundant pathways, where possible, are shown for each genotype. **(D)** Ingenuity pathway analysis was conducted on all significant DAR for Mtb-infected WT, ΔPPM1A and PPM1A^+^ macrophages. The radial summary graph of DAR in WT macrophages is shown. Radial summary graphs for ΔPPM1A and PPM1A^+^ macrophages are shown in **Figure S6**.

Significant DAR from each genotype were split by whether chromatin accessibility increased or decreased upon Mtb infection. GREAT was used to determine the biological pathways that were enriched in each group to associate biological functions to DAR where chromatin accessibility exclusively increased or decreased upon Mtb infection. In all genotypes, type I IFN responses was the most highly enriched GO biological processes in DAR where chromatin accessibility increased. However, enrichment of this term in PPM1A^+^ cells was 10 to 15-fold less than in WT or ΔPPM1A cells. In WT cells, the GO terms “PERK-mediated unfolded protein response” and “Induction of positive chemotaxis” were also highly enriched in DAR where chromatin accessibility increased, while the “L-serine metabolic process” was highly enriched in ΔPPM1A cells (**Fig. 2C**). In DAR where chromatin accessibility decreased, “Neural nucleus development” was the only GO term that was significantly enriched in any genotype, showing up in the PPM1A^+^ cells (**Fig. 2C**). Overall, pathway analysis indicated that regions of altered chromatin accessibility induced by Mtb infection are enriched for type I IFN signaling pathway genes. In addition, we used QIAGEN Ingenuity Pathway Analysis (IPA) (Krämer *et al*. 2014) to analyze all DAR between Mtb-infected and non-infected macrophages. The interaction network that was derived from DAR in WT cells identified the type I IFN transcriptional regulator IRF7 as the central hub, as well as several other genes involved in type I IFN responses such as *IRF3, IRF4, IRF5, DDX58/RIGI1* and *STAT2* (**Fig. 2D)**. Radial summary graphs from the IPA for ΔPPM1A and PPM1A^+^ macrophages are shown in Figure S4.

### Chromatin remodeling during Mtb infection may alter transcription factor binding

Using the ATAC-seq dataset, we performed transcription factor binding motif (TFBM) analysis to identify the transcription factors (TF) for which DNA accessibility is modulated at their predicted binding sites in WT, ΔPPM1A and PPM1A^+^ cells upon Mtb infection. In Mtb-infected WT cells, the ZNF460 binding site was the most abundant, and was found in 95 (34%) DAR where chromatin accessibility increased (**Fig. 3A)**. The enrichment of the ZNF460 binding motif was also present in Mtb-infected ΔPPM1A and PPM1A^+^ macrophages but were not statistically significant. Instead, the most abundant TF binding motif in ΔPPM1A cells was the ZNF143 binding site, which was found in 41 (17%) DAR where CA decreased (**Fig. 3B)**. Interestingly, both ZNF460 and ZNF143 are zinc finger TFs, but their differential enrichment in DAR where CA increased (WT) or decreased (ΔPPM1A) may highlight key functional differences in these two genotypes. The most enriched TF binding site in PPM1A^+^ cells was JDP2, which was found in 2878 (36%) DAR where CA increased (**Fig. 3A)**.

**Figure 3.**
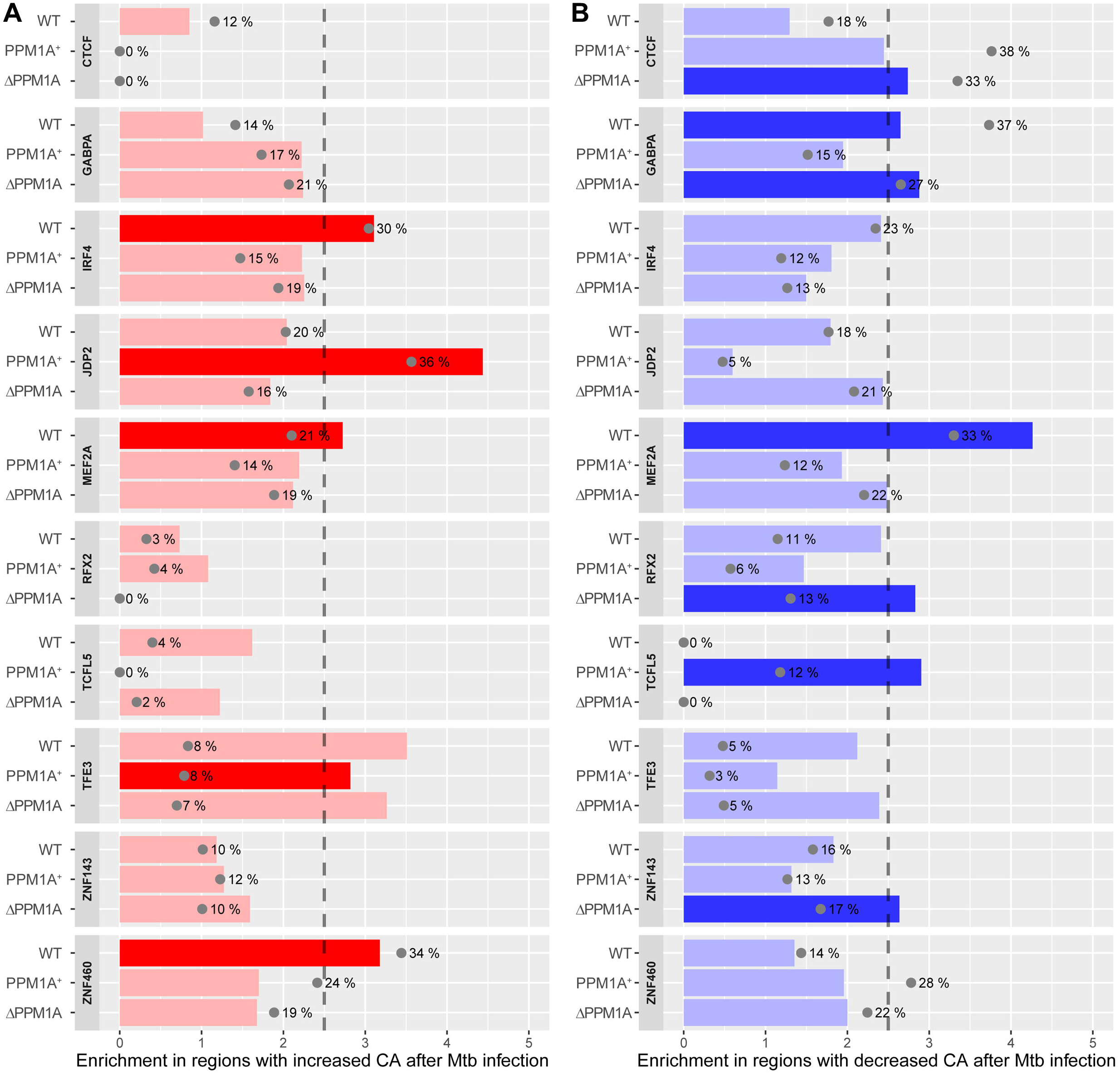
Transcription factor binding motif enrichment in Mtb-infected macrophages. TF binding motifs across all genotypes that were significantly enriched in ATAC-seq peaks with increased or decreased CA were identified; those with enrichment score above 2.5 and FDR < 0.001 were considered significant. **(A)** Enrichment and significance of the selected motifs in DAR with increased CA. Grey dots indicate the percentage of regions that have at least one hit to the motif. Bright red indicates motifs that are statistically significant. **(B)** Enrichment for the same motifs as shown in panel A, but within DAR with decreased CA following Mtb infection. Bright blue indicates motifs that are statistically significant.

TF motif enrichment analysis revealed that 5 TF binding motifs were significantly enriched following Mtb infection in WT cells, 4 in ΔPPM1A cells, and 3 in PPM1A^+^ cells **(Fig. S5)**. In DAR where chromatin accessibility increased, TF binding motifs for IRF4 (regulator of M2 polarization), ZNF460, and MEF2A were significantly enriched in WT cells, while binding motifs for JDP2 and TFE3 were enriched in in PPM1A^+^ cells, and there were no enriched TF binding motifs in ΔPPM1A cells **(Fig. 3A)**. Interestingly, TFE3 regulates autophagy and is normally shown to be repressed during Mtb infection (Pastore *et al*. 2016; Brady, Martina and Puertollano 2018). In DAR where chromatin accessibility decreased, TF binding motifs for GABPA and MEF2A, a regulator of type I IFN responses (Cilenti *et al*. 2021) in macrophages, were enriched in WT cells, while CTCF, GABPA, RFX2 and ZNF143 were enriched in ΔPPM1A cells and TCFL5 was enriched in PPM1A^+^ cells **(Fig. 3B)**. Other than GABPA, significant enrichment for all other binding motifs was unique to the genotype. Overall, motifs that were highly enriched in a specific genotype was also specific to the change in chromatin accessibility (*i.e*. increased or decreased CA), which support the TFBM analysis. The data also show that PPM1A expression modulated the TFBM that were enriched at DAR following Mtb infection. For example, deletion of PPM1A enriched for TFBM (4 motifs) specifically in regions where CA decreased, while overexpression of PPM1A resulted in more and stronger TFBM enrichment (2 motifs) in regions where CA increased.

### Mtb infection alters transcriptome in WT, ΔPPM1A and PPM1A^+^ macrophages

We performed RNA-seq in parallel to ATAC-seq to correlate changes to gene expression with the DAR induced by Mtb infection. There were numerous changes to global gene expression in Mtb-infected WT, ΔPPM1A and PPM1A^+^ cells compared to non-infected cells (**Fig. 4A, B**). As observed with DAR, there was also a significant increase in the total number of differentially expressed genes (DEG) in PPM1A^+^ cells compared to WT and ΔPPM1A cells (**Fig. 4A, B**). Using edgeR, we identified that Mtb infection induced 410 significant DEG (298 up and 112 downregulated) in WT cells, 508 significant DEG (338 up and 170 downregulated) in PPM1A cells, and 936 significant DEG (713 up and 223 downregulated) in PPM1A^+^ cells **(Fig. 4C**). While PPM1A^+^ cells had an approximately two-fold increase in total DEG compared to WT and ΔPPM1A cells, this proportion is considerably less than the difference in total DAR between PPM1A^+^ cells and WT (22-fold) or ΔPPM1A (15-fold) cells. We identified 33, 94 and 456 unique significant DEG upon Mtb infection in WT, ΔPPM1A and PPM1A^+^ cells, respectively **(Fig. 4C, D)**. In WT and ΔPPM1A cells, there were more unique DEG that were downregulated (21 and 61, respectively) than upregulated (12 and 33). In contrast, there were more unique DEG that were upregulated (337) than downregulated (119) in PPM1A^+^ cells **(Fig. 4C**). There was 326 DEG shared between WT and PPM1A, 298 between WT and PPM1A^+^ cells, 330 between ΔPPM1A and PPM1A^+^ cells, and 258 DEG shared between all genotypes **(Fig. 4D)**.

**Figure 4.**
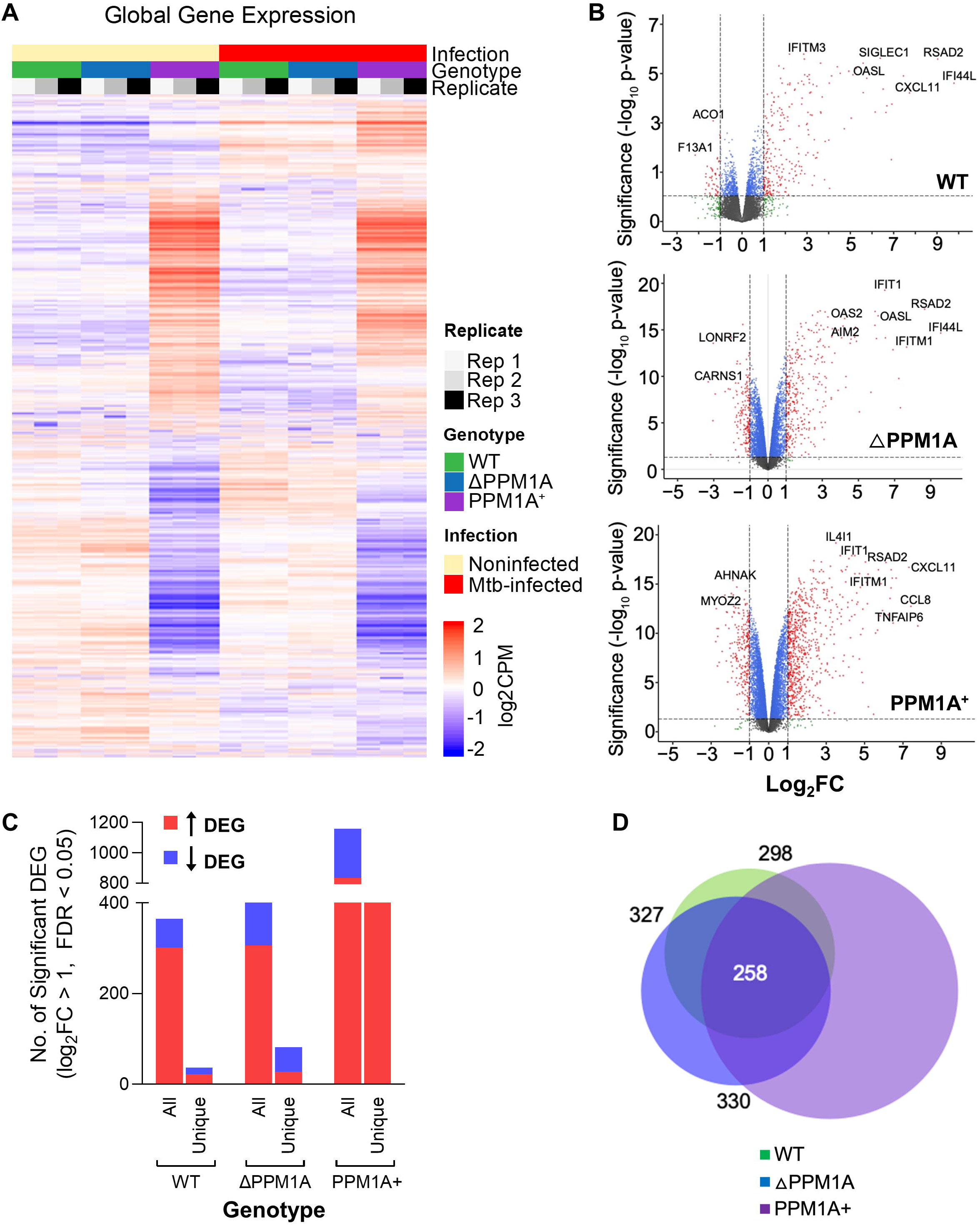
Mtb infection alters gene expression in WT, ΔPPM1A and PPM1A^+^ macrophages. **(A)** Global gene expression. Heatmap of global gene expression in WT, ΔPPM1A and PPM1A^+^ cells before and after Mtb infection are shown. Genes with log_2_CPM > 0 counts were considered significantly expressed. **(B)** Volcano plots displaying differentially expressed genes (DEG) in WT, ΔPPM1A and PPM1A^+^ macrophages upon Mtb infection. DEG with absolute value of Log2FC > 1 and FDR < 0.05 were considered significant. Gene names of the most striking up and downregulated genes are shown. **(C)** Graph showing the total and unique numbers of DEG in each genotype. Red or blue bars show the number of DEG that increased or decreased in expression, respectively. **(D)** Venn diagram showing the overlap of significant DEG between Mtb-infected WT, ΔPPM1A and PPM1A^+^ macrophages and non-infected controls.

Notable DEG unique to WT cells included upregulated *C1QA*, which is a strong marker of active TB (Dijkman *et al*. 2020), and downregulated *NOXA*, which is involved with nitric oxide mechanisms. DEG unique to ΔPPM1A cells include *IL1A*, which is important for Mtb clearance, T cell priming, and granuloma formation (Di Paolo *et al*. 2015; Lovey *et al*. 2022). In PPM1A^+^ cells, several interesting genes are upregulated, including *IL24* and *IL32*, which are known as correlates of protection against TB (Ma *et al*. 2011; Montoya *et al*. 2014; Koeken *et al*. 2020). PCA analysis of total gene expression data also displayed that the largest contributor to variance in the dataset was genotype. Replicates clustered together based on genotype and infection, confirming the changes to gene expression that we observed **(Fig. S6).**

### Gene expression profiles of Mtb-infected macrophages are enriched in type I IFN responses and anti-viral signaling pathways

Gene set enrichment analysis (GSEA) was conducted to identify gene sets that were enriched in significant DEG in WT, ΔPPM1A and PPM1A^+^ macrophages upon Mtb infection compared to non-infected cells (Mootha *et al*. 2003; Subramanian *et al*. 2005). We focused the GSEA analysis only on GO biological processes to enable a direct comparison to pathway analysis generated by GREAT in the ATAC-seq pipeline **(Fig. 2)**. The most highly enriched GO biological processes in significant DEG in cells upon Mtb infection were “Defense response to virus”, “Response to type I IFN”, “Negative regulation of viral processes” and “Response to IFN-γ” **(Fig. 5A)**. The same GO biological processes were highly enriched in ΔPPM1A and PPM1A^+^ cells, as well as “Granulocyte chemotaxis”. Interestingly, the most highly enriched GO biological processes in both ΔPPM1A and PPM1A^+^ cells was “Co-translational protein targeting to membrane”, which along with “Establishment of protein localization to the endoplasmic reticulum”, was enriched in downregulated genes **(Fig. 5A)**. There were more common GO terms between ΔPPM1A cells and PPM1A^+^ cells genotypes (**Fig. 5B)**. GSEA plots generated for the GO biological process terms “Response to type I IFN”, and “Defense Response to Virus” denoted the leading-edge genes that most influenced the enrichment of these terms **(Fig. S7A-C)**. The median-centered log_2_FC was greater for leading-edge genes in PPM1A^+^ cells than WT or ΔPPM1A cells upon Mtb infection (**Fig. 5C)**. Notably, the expression of several leading-edge genes from these GO terms, such as *MMP12, ISG20, WNT5A* and *HLA-F* (type I IFN signaling), *BIRC3, ZC3H12A, TLR8, DDX60 and STAT2* (Defense response to virus) changed in opposite directions in PPM1A^+^ cells compared to WT cells upon Mtb infection. Similarly, in PPM1A^+^ cells compared to ΔPPM1A cells, the expression of the leading-edge genes including *ERG1, HLA-E/B, IRF2, TRIM56, MX2, NLRC5, OASL, OAS2, IFI16* and *APOBEC3F/G* (Type I IFN signaling), and *NT5C3A, MLKL* and *NLRC5* and *IFI6* (Defense response to virus) changed in the opposite direction upon Mtb infection. Additionally, all DEG sets between Mtb-infected and non-infected cells were analyzed with QIAGEN IPA. The interaction network that was derived from the significant DEG upon Mtb infection in WT cells identified the type I IFN transcriptional regulator IRF7 as the central hub, as well as several other genes involved in type I IFN responses such as *IRF3, TLR3, TLR9, DDX58/RIGI1* and *STAT1* (**Fig. 5D)**. Radial summary graphs from the IPA for ΔPPM1A and PPM1A^+^ macrophages are shown in **Figure S8**.

**Figure 5.**
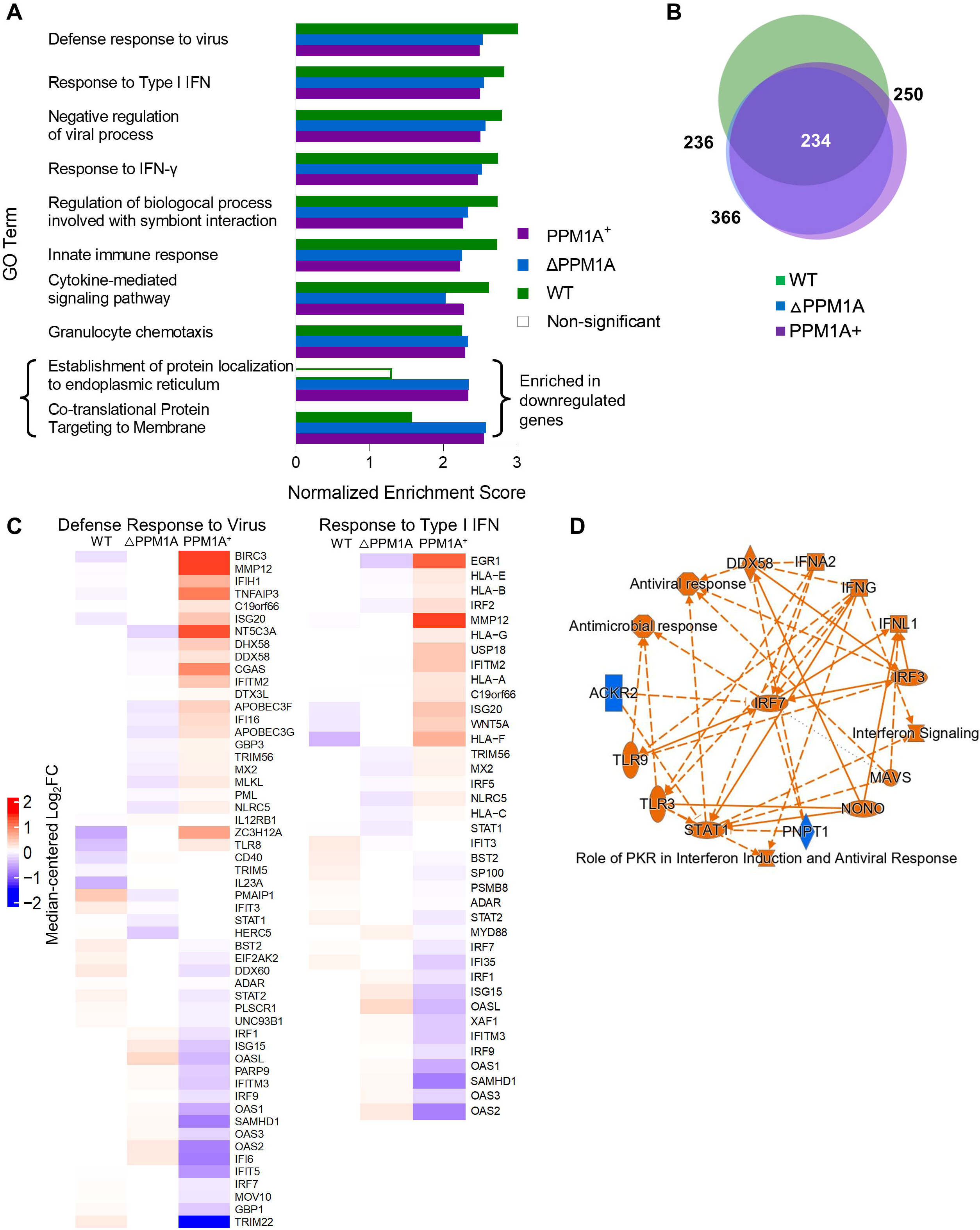
Gene Set Enrichment Analysis (GSEA) identifies a prominent type I IFN signatures in Mtb-infected macrophages. **(A)** GSEA was conducted to determine gene sets that were enriched in WT, ΔPPM1A and PPM1A^+^ macrophages upon Mtb infection. Results that were GO biological process terms were selected for analysis. GO terms with FDR < 0.05 and a set size of at least 10 genes were considered significant. GO terms were ranked by the absolute value of the normalized enrichment score (NES) and the top 5 non-redundant terms from each genotype were plotted by NES. **(B)** Venn diagram displaying common significant GO terms between WT, ΔPPM1A and PPM1A^+^ macrophages after Mtb infection. **(C)** Differentially expressed leading-edge genes. Two highly enriched GO biological process gene sets (“Response to Type I IFN” and “Defense Response to Virus”) are shown. The corresponding heatmaps display normalized log_2_FC differential expression values that were median-centred across all samples of the dataset. **(D)** Ingenuity pathway analysis was conducted on all significant DEG for Mtb-infected WT, ΔPPM1A and PPM1A^+^ macrophages. The radial summary graph of DEG in WT macrophages is shown. Radial summary graphs for ΔPPM1A and PPM1A^+^ macrophages are shown in **Figure S8**.

### Mtb infection triggers a type I IFN response that is concordant across chromatin accessibility and transcriptional signatures

We investigated the functional significance of changes to CA in Mtb-infected WT, ΔPPM1A and PPM1A^+^ cells compared to non-infected controls by evaluating correlating changes to gene expression in these regions. Overlapping DAR and DEG were determined by matching the gene IDs that were annotated to DAR with DEG. For genes with multiple CA peaks annotated to the same gene, the peak closest to the TSS of the gene was selected to be the match for further analysis. In WT cells, 34 genes were differentially expressed and located near one or more DAR, with 30 genes showing increased CA and gene expression, and 4 genes showing decreased CA and gene expression **(Fig. 6A & Fig. S9A)**. In ΔPPM1A cells, there were 44 genes that were differentially expressed and located near one or more DAR (10% of total DAR and 9% of total DEG), where CA and gene expression increased in 31 and 34 genes, and decreased in 13 and 10 genes, respectively **(Fig. 6A & Fig. S9A)**. In PPM1A^+^ cells, there was 347 overlapping genes (6% of total DAR and 37% of total DEG), where CA and gene expression increased in 283 and 266 genes, and decreased in 63 and 80 genes, respectively **(Fig. 6A & Fig. S9A)**. In all genotypes, the most significant overlapping genes, with the greatest changes to gene expression were all located in peaks where CA increased **(Fig. 6A & Fig. S9A)**. Notably, in each genotype, the most significant overlapping genes were predominantly type I IFN response genes, and were among the most significant DEG. In Figure 6B, we plotted significant overlapping genes by log_2_FC gene expression versus log_2_FC CA to determine the correlation of change in genes upon Mtb infection. In WT and ΔPPM1A cells, the log_2_FC of overlapping genes were highly positively correlated. In PPM1A^+^ cells, the correlation of change increased, but less so than in WT and ΔPPM1A due to lower maximum change in DEG, and an increased number of DAR and DEG that decrease (**Fig. 6B).** Notably, the majority of DAR that overlapped with DEG were annotated to introns in all genotypes **(Fig. S9B)**, compared to all DAR which were primarily annotated to “Other introns” and “Distal intergenic” regions **(Fig. S2F).** We compared the change in CA and gene expression of the top 10 most highly up and downregulated overlapping genes in each genotype, ranked by log_2_FC gene expression (**Fig. 6C**). In all genotypes, changes to gene expression were greater than changes to CA, except for *C1S*, *IFI44* and *IFIT1* in PPM1A^+^ cells. Furthermore, 3 overlapping genes are unique to WT cells, 16 to ΔPPM1A cells and 316 to PPM1A^+^ cells (highlighted).

**Figure 6.**
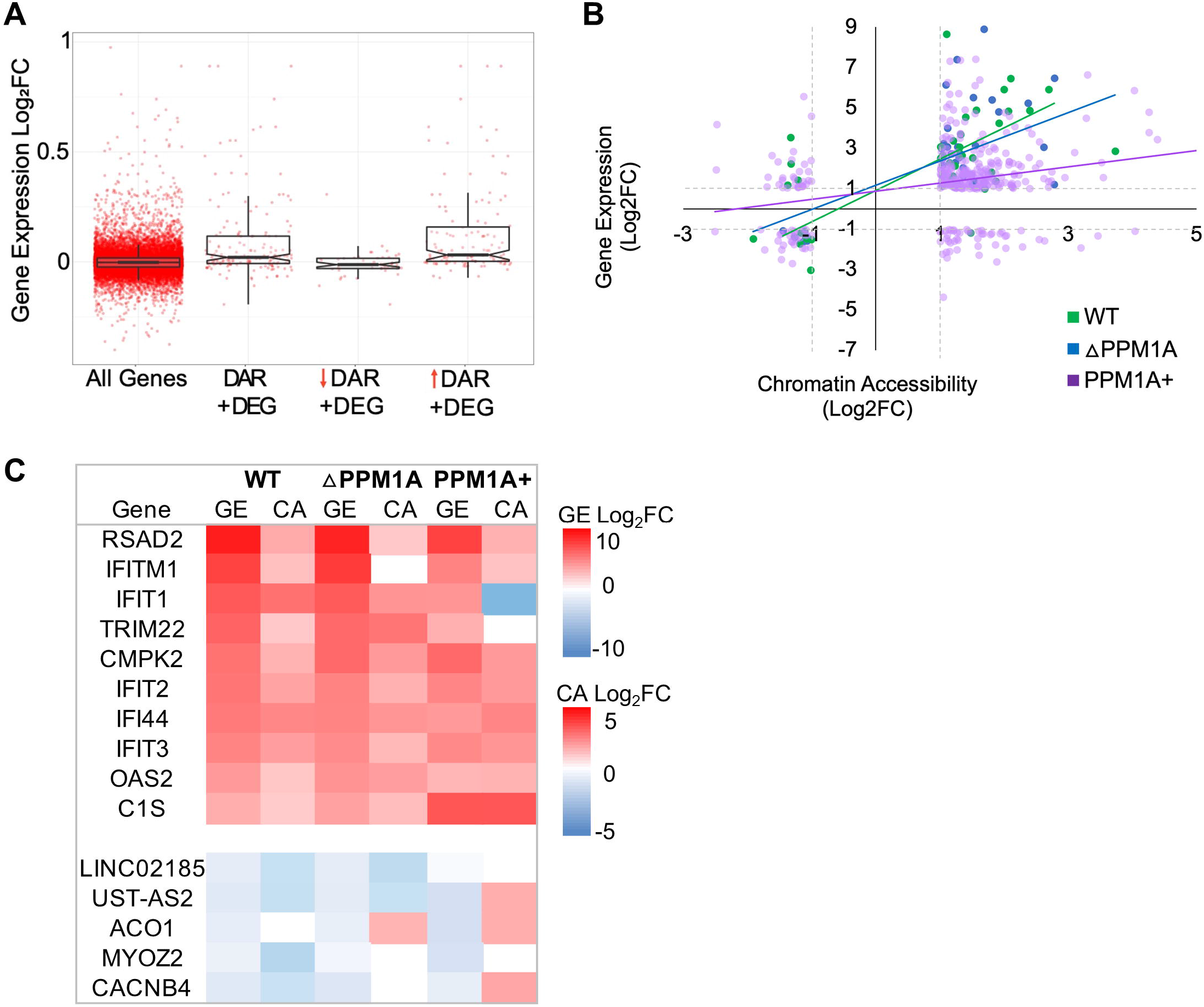
Differential chromatin accessibility is concordant with differential gene expression. **(A)** Boxplot of DEG that are concordant with DAR (DAR+DEG) in WT, ΔPPM1A and PPM1A^+^ macrophages upon Mtb infection. All DEG were matched to DAR based on the gene names that DAR were annotated to. In cases where multiple peaks annotated to the same gene, the peak closest to the TSS of the gene was used. DEG that are concordant with DAR were separated based increasing or decreasing CA. **(B)** Correlation of DAR and DEG. The log_2_FC of all DEG in each genotype was plotted against the log_2_FC of the matching DAR. **(C)** The top 10 highest DEG that are concordant with DAR in all genotypes upon Mtb infection were ranked by log_2_FC of gene expression. Log_2_FC values are displayed for each genotype, even if they were not significant. Heatmap colour scales represent the magnitude of change (Log_2_FC) in either DEG or DAR.

## DISCUSSION

This study showed that Mtb infection induces genome-wide chromatin remodeling and changes to gene expression. In addition, the expression of PPM1A significantly modulated the total number and location of chromatin regions where accessibility changed, and genes that were differentially expressed. Significant DAR and DEG were highly enriched in type I IFN responses. Type I IFN enrichment in DAR is largely driven by regions where CA increased. Furthermore, we identified subsets of type I IFN response genes in each genotype with striking and concordant changes to chromatin accessibility and gene expression. These genes partially overlapped between genotypes, but certain genes were unique to WT, ΔPPM1A, or PPM1A^+^ cells. Many TF motifs that were enriched in each genotype were motifs for TF associated with TB pathogenesis or known to regulate monocyte differentiation. PPM1A expression also modulated the TF motifs that were enriched, as well as whether certain motifs were enriched in DAR where CA increased or decreased.

The only previous report to investigate global CA patterns upon Mtb infection was a report by Correa-Macedo et al. where the authors investigated CA and gene expression patterns of primary human alveolar macrophages (hAM) infected with Mtb *ex vivo* (Correa-Macedo *et al*. 2021). They report over 12,360 DAR and 1434 DEG total in hAM upon Mtb infection, without applying any log_2_FC cut-offs. Applying abs(Log_2_FC > 1) and FDR < 0.05 cutoffs to their data reduces the number of DAR to six, where CA increases, and the number of DEG to 126, where 123 are upregulated and 3 are downregulated. Considering that they infected with MOI 5, the parallel number of DAR and DEG is still significantly less than reported in our study. Given that multiple studies in addition to ours report between 600 and 3000 DEG in macrophages upon Mtb infection using these standard cutoffs (Wu *et al*. 2012; Seshadri *et al*. 2017; Papp *et al*. 2018; Roy *et al*. 2018; Lee *et al*. 2019; Looney *et al*. 2021; Pu *et al*. 2021), it appears as though there may be a limiting factor in the data reported by Correa-Macedo et al.. This may be due to the phenotypic and functional differences between hAM and THP-1 cells, conditions of Mtb infection (MOI, time, strain), or variability between patient samples caused by increased genomic heterogeneity. Interestingly, the DAR that Correa-Macedo et al. reported are annotated to many of the same genes that the DAR in our analysis annotated to (*RSAD2, MX1,2, IFIT1,2,3*, and *IFITM3*) Pathway analysis by Correa-Macedo et al. also demonstrated enrichment of type I IFN signalling pathways, while IRF9 and ZNF685 motifs, which are involved with type I IFN signaling responses, were significantly enriched in TFBM analysis (Correa-Macedo *et al*. 2021). They also report changes to CA at the TSS of *CXCL10, IFI44L, APOBEC3A*, and *MX1*, which are among genes with highest log_2_FC identified in our DAR. In our integrative analysis, *MX1* is significant in WT cells, but *CXCL10, IFI44L* and *MX1* are significant in PPM1A^+^ cells. Correa-Macedo et al. provide a foundational and clinically relevant investigation of CA patterns in Mtb infected hAM. However, their focus was on CA and gene expression patterns of HIV patients and patients receiving pre-exposure prophylaxis therapy, rather than Mtb infection, so an in-depth investigation of CA pathways in Mtb infection alone was not the priority of their study.

We determined that there were significantly more genes that were upregulated than downregulated upon Mtb infection in macrophages. This is consistent with previous transcriptome analyses of Mtb infected macrophages (Wu *et al*. 2012; Seshadri *et al*. 2017; Papp *et al*. 2018; Roy *et al*. 2018; Lee *et al*. 2019; Looney *et al*. 2021; Pu *et al*. 2021). We also found that there was almost an equal number of DAR where CA increased or decreased upon Mtb infection in macrophages. This is consistent with previous reports that also demonstrate increases in host CA upon Mtb infection, but DAR where host CA decreases is seldom reported (10–15, 24). One report demonstrated CA decreasing in macrophages at the *IL12B* locus due to Mtb-induced upregulation of HDAC1, which resulted in impaired autophagy (Chandran *et al*. 2015). Limited data on decreasing CA upon Mtb infection could be due to the ChIP-seq methods used in these studies, which indirectly infer CA by the location of a specific histone mark throughout the genome, thereby limiting these analyses to the probe used. Indeed, decreasing CA and gene repression, is the target of many secreted Mtb effector proteins that directly induce histone modifications (Gauba *et al*. 2021). Therefore, in the present study, we have captured a more complete representation of global changes to CA by evaluating both increases and decreases to CA.

In both CA and transcriptome analyses, PPM1A overexpression contributes to large-scale chromatin-remodeling to increase accessibility and induce transcription. As PPM1A is a phosphatase, this suggests that histone-modifying enzymes may be activated or repressed by dephosphorylation to mediate opening of chromatin upon Mtb infection. The unusually large number of DAR in the PPM1A^+^ cells may be partially a result of artificially high overexpression of PPM1A (Berton *et al*. 2022a), which may induce global non-specific dephosphorylation effects or compensatory effects in the cell line due to the broad number of pathways influenced by PPM1A activity (Li *et al*. 2015, 2022; Berton *et al*. 2022b). We also observe this phenotype through differences in baseline CA and gene expression, compared to WT cells, (**Fig. 1C & Fig. 4A, B**), and by PCA (**Fig. 1D & S6**). Interestingly, despite that PPM1A^+^ cells have a ~20-fold increase in the number of CA compared to WT cells, this only led to a modest ~2-fold increase in the number of DEGs. This indicates that cells have strict control at various levels of processes leading to transcription and protein translation that can be compensated, consistent with a recent report of protein dosage compensation (Schukken and Sheltzer 2022). However, the subset of DAR and DEG that are unique to ΔPPM1A cells compared to WT and PPM1A^+^ cells indicate that PPM1A activity is also involved in gene repression. This is the mechanism illustrated in *Listeria monocytogenes* infection where PPM1A dephosphorylates the histone deacetylase SIRT2 to induce chromatin remodeling and gene repression, which is required for infection (Pereira *et al*. 2018: 2). While PPM1A could impact CA directly, by dephosphorylating chromatin-binding proteins or modulating the activity of a chromatin-modifying enzyme such as SIRT2, we cannot rule out that it might additionally affect CA indirectly, by influencing the expression levels of genes that participate in regulating CA. Nevertheless, we showed that Mtb infection induces changes in WT, ΔPPM1A and PPM1A^+^ macrophages that are predominantly enriched in type I IFN GO pathways for both DAR and DEG.

The type I IFN response triggered by Mtb infection in macrophages and as a transcriptional signature of TB disease is well characterized (Stanley *et al*. 2007; Dorhoi *et al*. 2014; Ji *et al*. 2019). Numerous transcriptome analyses provide evidence that type I IFN response genes are enriched among upregulated genes in Mtb-infected macrophages (Stanley *et al*. 2007; Berry *et al*. 2010; Singhania *et al*. 2018; Lavalett, Ortega and Barrera 2020). There is a large body of evidence supporting that upregulated type I IFN responses drive host pathology in TB and enhance Mtb survival in macrophages. For example, type I IFN responses impair host-protective IFN-γ and IL-1 signalling, induce pathogenic IL-1Ra expression, promote early cell death in alveolar macrophages (AM), and promote neutrophil-induced pathology (Park *et al*. 2017). Studies also show that the immune response in TB is more effective when the IFN-α receptor is blocked to limit the type I IFN response (Manca *et al*. 2005; Ordway *et al*. 2007; Stanley *et al*. 2007; Mayer-Barber *et al*. 2011; Dorhoi *et al*. 2014; Kimmey *et al*. 2017). Importantly, this is also demonstrated in AM from TB patients (Lavalett, Ortega and Barrera 2020; Correa-Macedo *et al*. 2021) and murine AM where it was associated with impaired metabolism and mitochondrial stress (Olson *et al*. 2021). In agreement with this knowledge of pathogenesis and immunity in TB, we showed that significant DAR and DEG in macrophages are enriched in type I IFN response pathways. Pathways such as ‘Type I IFN signalling pathway’, ‘Defense response to virus and ‘Response to IFN-α’ were more enriched in DAR where CA increased, indicating that upregulation of genes in these pathways that we and others observe, occurs at the level of CA (**Fig. 2C and Fig. 5A**). We uncovered a small subset of genes in type I IFN response pathways that were both the most significant and highly changed DAR and DEG (**Fig. 6C**), suggesting that these genes are regulated at the level of CA in macrophages upon Mtb infection. Furthermore, changes in CA and gene expression in these genes were highly correlated (**Fig. 6C**). There were genes within this subset that were unique to either WT, ΔPPM1A or PPM1A^+^ cells, suggesting that PPM1A modulates CA and gene expression even among this small gene subset (**Fig. 6F**).

Interestingly, type I IFN are also involved in chromatin remodeling and trained immunity in macrophages. IFN-α stimulation in concert with TNF activity was observed to “prime” chromatin by enhancing tri-methylation of H3K4, which blocked downregulation of inflammatory genes and enhanced their responsiveness (Park *et al*. 2017). Macrophage stimulation with IFN-β has also been shown to remodel 3D chromatin structure and increases CA at many interferon-stimulated genes. Notably, this effect was prominent at *Oas, Mx1,2* and *Ifit1* genes (Platanitis *et al*. 2022), which are among the subset of genes we show to be DAR and DEG upon Mtb infection. In fibroblasts, stimulation with IFN-β also leads to accumulation of H3K36me3 at *Oas1, Mx1* and *Ifit1*, termed “memory interferon-stimulated genes”, mediating enhanced upregulation upon re-stimulation (Kamada *et al*. 2018). Impairing innate immune memory in myeloid progenitors is also an evasion strategy of Mtb that is mediated through type I IFN signaling, which leads to impaired iron metabolism, mitochondrial function and induced necrosis in myeloid progenitors from Mtb-infected mice (Khan *et al*. 2020). Our data showing that induction of type I IFN genes occurs at the level of CA upon Mtb infection supports this unique function of type I IFN as modulators of chromatin structure and trained immunity. Manipulating trained immunity as an evasion strategy benefits Mtb by re-programming multiple generations of cells. As such, knowledge on specific genes that can act as markers for modified CA during Mtb infection could prove to have prognostic or therapeutic value.

Transcription factor binding motif analysis of ATAC-seq data suggests that MEF2A (or another TF with similar DNA sequence binding preference, such as MEF2B, C or D) activity is modulated upon Mtb infection in WT macrophages (**Fig. 3**). MEF2A is commonly studied in the context of cardiac muscle cell development (Ornatsky and McDermott 1996; Wales *et al*. 2014; Xiong *et al*. 2019), angiogenesis (Sacilotto *et al*. 2016), and recently was shown to critically regulate expression of type I IFN genes in macrophages (Cilenti *et al*. 2021). MEF2A regulated genes included *IFIT1, RSAD2, OASL2, MX1* and *IRF7*, which we show in Mtb infection is mediated by increasing CA. As such, type I IFN responses induced by Mtb infection could be mediated by increased or decreased activity of genes regulated by MEF2A given that its binding motifs are enriched in both DAR where CA increases and decreases **(Fig. 3)**. Towards this, MEF2A reportedly upregulates PIK3CG (Xiong *et al*. 2019), which inhibits IL-17A secretion, and is downregulated in advanced TB patients (Leisching 2018). Importantly, MEF2A-PIK3CG signaling was reported to increase SIRT1 activity (Liu *et al*. 2019). SIRT1 is a histone deacetylase that is downregulated upon Mtb infection to repress autophagy, phagosome-lysosome fusion and promote pathology (Cheng *et al*. 2017). Thus, dysregulated MEF2A activity observed upon Mtb infection may be a link to SIRT1 downregulation. In addition, IRF4, a less well characterized TF of the IRF family, was highly enriched in WT macrophages upon Mtb infection. IRF4 plays an important role in regulating myeloid cell development, differentiation, and inflammation, as it binds to MyD88 and promotes M2 polarization (Jefferies 2019). However, IRF4 has not been studied in the context of Mtb infection.

Our TFBM analysis also revealed that JDP2 activity is modulated by PPM1A expression upon Mtb infection (**Fig. 3**). It is however important to note that multiple other related family members could bind to the same motifs. JDP2 is member of the large AP-1 family of TF, but acts as a repressor of AP-1 TF, such as c-JUN and ATF2, which are repressed through recruitment of HDAC3 (Jin *et al*. 2006; Nakade *et al*. 2007). JDP2 has multiple roles in epigenetic modulation as it reportedly binds to histone 3 and 4, has histone chaperone activity associated with nucleosome disassembly and inhibits the histone acetyltransferase p300 (Jin *et al*. 2006; Nakade *et al*. 2007). JDP2 is phosphorylated by JNK, which interestingly is inactivated by PPM1A dephosphorylation (Katz, Heinrich and Aronheim 2001), suggesting that JNK could link PPM1A and JDP2 activity. This corroborates our findings that JDP2 binding sites are enriched in PPM1A^+^ cells, and that this pathway is enriched upon Mtb infection. The AP-1 family member ATF3 is a negative regulator of IFN-β in macrophages, which aligns with our finding that type IFN responses are less enriched in PPM1A^+^ cells where JDP2 motifs are highly enriched (Labzin *et al*. 2015). A third AP-1 family member with potential to bind the enriched JDP2 motif, BATF2, is upregulated in pro-inflammatory (M1) macrophages upon Mtb infection, and induces the expression of several key inflammatory mediators (Roy *et al*. 2015). The incredibly diverse role of AP-1 TF in macrophages upon infection poses a challenge to directly link enrichment with specific aspects of PPM1A overexpression. Nonetheless, enrichment of JDP2 binding motif in PPM1A^+^ cells provide evidence that PPM1A activity modulates key inflammatory responses in concert with TF that can change the course of anti-bacterial immune responses. Interestingly, both MEF2 and ATF family TF binding sites are enriched in enhancers found within Alu repeats in macrophages. Genes associated to these enhancer regions specifically regulate lipid cholesterol metabolism in macrophages upon Mtb infection (Bouttier *et al*. 2016)

A limitation of this study is that we broadly analyzed CA based on chromatin structure, when it is regulated by many factors including DNA methylation, miRNA and lncRNA. Therefore, the changes in CA that we have identified may be mediated by another, or likely multiple mechanisms. To account for this, datasets for DNA methylome and miRNA-ome of Mtb infection could be cross-referenced to extend the mechanistic basis of our conclusions. Additionally, methylation specific-PCR could be performed to determine the methylation status of the identified genomic regions of interest. Further analysis of the 3D structure of DNA could be investigated using HiChIP. It is also important to note that this study was conducted exclusively with THP-1 macrophages, and while many identified signatures were consistent with literature from primary human monocyte derived macrophages, it would be prudent to confirm key findings in primary human macrophages, including alveolar macrophages, which are the most clinically relevant model for Mtb infection (Madden *et al*. 2022).

This study reveals that significant genome-wide changes to CA are induced by Mtb infection of macrophages, and that expression of a subset of type I IFN response genes is mediated by these changes. Our results have implications for TB research because of the major role of type I IFN responses in disabling the human immune response to Mtb infection. This provides a novel way that the identified genes and pathways could be developed as biomarkers or targeted for adjunctive therapy. With the rise in cases of multi-drug resistant TB and lack of novel antibacterial therapies, developing potent adjunctive therapy is imperative to combatting the world’s deadliest bacterial pathogen.

## MATERIALS AND METHODS

### Cell culture

THP-1 monocytes (ATCC TIB-202) were maintained in RPMI 1640 medium (Gibco, Gaithersburg, MD). RPMI 1640 was supplemented with 2 mM L-glutamine, 100 I.U./ml penicillin, 100 μg/ml streptomycin, 10 mM HEPES, and 10% heat-inactivated fetal bovine serum purchased from Gibco. THP-PPM1A^+^ and THP-ΔPPM1A cells were generated previously as described (Sun *et al*. 2016; Berton *et al*. 2022a). Cells were maintained at 37°C in a humidified atmosphere of 5% CO_2_. THP-1 monocytes were differentiated with 100 ng/ml phorbol ester 13-phorbol-12-myristate acetate (PMA, Alfa Aesar, Haverhill, MA) for 72 h.

### Bacterial strains and infection

The *Mycobacterium tuberculosis* H37Rv-derived auxotroph strain mc^2^6206 (Sampson *et al*. 2004; Mouton *et al*. 2019) was grown in Middlebrook 7H9 medium (BD Biosciences, Franklin Lakes, NJ) supplemented with 0.2% glycerol (Fisher Chemical, Waltham, MA), 0.05% Tween-80 (Acros Organics, Fair Lawn, NJ), 10% OADC (BD Biosciences), 24 μg/ml D-pantothenic acid (Alfa Aesar), and 50 μg/ml L-leucine (Alfa Aesar). Liquid Mtb cultures were maintained at 37°C with slow shaking (50 rpm).

Mtb mc^2^6206 growing in log-phase was quantified by optical density measurement at 600 nm using the conversion of OD 1 = 3 × 10^8^ Mtb bacteria per ml. The number of bacteria required for a multiplicity of infection (MOI) of 10 was washed and resuspended in RPMI 1640 cell culture media without antibiotics. Bacteria were added to the differentiated THP-WT, THP-ΔPPM1A, or THP-PPM1A^+^ macrophages and incubated at 37°C for 16 h. Three phosphate buffered saline (PBS) washes were then performed to remove extracellular, non-phagocytosed bacteria and infection was continued at 37°C for a total infection time of 48 h. Non-infected control cells underwent identical wash and incubation steps.

### ATAC-seq library preparation

ATAC-seq of infected and non-infected macrophages was performed according to published OMNI-ATAC protocols (Buenrostro *et al*. 2013; Corces *et al*. 2017). Tn5 transposomes were generated following the protocol of Picelli et al., using pTXB1-Tn5 (gift from Rickard Sandberg, (Addgene plasmid # 60240)) (Picelli *et al*. 2014: 5). Transposomes were used with 2X Tagmentation DNA buffer (20 mM Tris-HCl, 10 mM MgCl_2_, 20% dimethyl formamide) from the OMNI-ATAC protocol. Macrophages were washed with PBS and harvested from cell culture plates using TrypLE Express Enzyme (Gibco). Fifty thousand Mtb-infected or non-infected THP-WT, THP-ΔPPM1A or THP-PPM1A^+^ cells were processed for ATAC-seq in duplicate. Each transposition reaction was cleaned up individually using Qiagen MinElute PCR purification kits. Transposed DNA libraries were amplified for 4-8 cycles using NEBNext High-Fidelity 2X PCR Master Mix. Individual DNA libraries were amplified using unique dual barcoding primers with unique 5’ i5 and 3’ i7 barcode sequences between the flow cell binding sequence and sequencing primer start site. PCR amplification reactions were purified using MagBio HighPrep PCR clean up kits to remove DNA fragments less than 50 bp and greater than 1000 bp. Pooled DNA libraries were sequenced in one lane of an S1 flowcell on the Illumina NovaSeq 6000 platform at The Centre for Applied Genomics, The Hospital for Sick Children, Toronto, Canada. Libraries were sequenced using 50 bp paired-end reads to reach a target of 30-50 million reads per sample. The DNA fragments tagmented by the Tn5 transposase were enriched in loci that are accessible (*RPLP0, PRKAB1*) in WT THP-1 macrophages under homeostasis, compared to those that are inaccessible (*CHRNA1, NANOG*) (30, 31).

### Raw data processing

Fasta sequence files were analyzed for quality using fastQC v0.11.9 (*FastQC* 2015). Remaining adaptor sequences and low-quality sequence fragments were removed using fastp v0.20.1 with the parameters --length_required 25, --cut_tail, --cut_tail_window_size 4, -- cut_tail_mean_quality 20, --disable_quality_filtering (Chen *et al*. 2018). Filtered reads were aligned to the human hg38 genome using STAR v2.7.5a without gene annotation GTF file, with the parameters -- alignIntronMax 1 --alignEndsType Extend5pOfReads12 --alignMatesGapMax 2000, --outFilterMatchNminOverLread 0.50, --outFilterScoreMinOverLread 0.50 and filtering for properly paired mates with a mapq score of 40 or above (Dobin *et al*. 2013) and duplicates were marked using Picard (Picard Tools). deepTools *AlignmentSieve* with parameter --ATACshift was used to adjust sequencing pairs by +5 and −4 base pairs to account for the 9bp shift introduced at Tn5 insert sites (Ramírez *et al*. 2016). Genome-wide sequencing coverage was calculated using deepTools v3.5.0 *bamCoverage* function and then converted to bigWig tracks that were normalized by counts per million (CPM) in each sample. ATAC-seq peak calling was conducted separately on each replicate using MACS2 with q-value of 0.00001 cutoffs, and peaks within blacklisted regions were removed with BEDTools (Gaspar 2018: 2). We used rmspc to identify consensus peaks which we defined at peaks that were found in at least two samples (Jalili *et al*. 2015; Jalili, Cremona and Paluzzi). Therefore, peaks that were present only in two replicates of one condition would be retained with parameters -r bio -w 1e-5 -s 1e-10 -c 2 and using the p-value calculated by MACS2 as the ‘value’.

### Differential accessibility analysis

Differential peak analysis was performed to compare significant differentially accessible chromatin regions in Mtb-infected macrophages versus non-infected macrophages. To qualify for differential accessibility testing, all peaks were required to have at least 20 Tn5 insertions sites in two samples. Qualifying peaks were analyzed with trimmed mean of M-values (TMM) normalization and batch effects were removed with RUVseq using the RUVs algorithm. Empirical tests determined that using a k value of 2 was sufficient to remove unwanted variation (Risso *et al*. 2014; Peixoto *et al*. 2015). Differential peak testing was then performed with edgeR and glmQLFTest function to contrast Mtb-infected cells of each genotype with corresponding non-infected cells (Robinson, McCarthy and Smyth 2010). Differentially accessible peaks with Benjamini-Hochberg adjusted P value (FDR) less than 0.05 and absolute log_2_FC value of at least 1 were considered significant. Heatmaps were generated using the ComplexHeatmap package (Gu, Eils and Schlesner 2016). Overlapping peaks between each genotype were calculated and plotted with the BioVenn R package (Hulsen, de Vlieg and Alkema 2008).

### Peak annotation and pathway analysis

Significant differentially accessible peaks were annotated to the human hg38 genome using ChIPseeker to determine the genes or regulatory elements in proximity to the genomic locations of peaks. Peaks were annotated to transcription start sites (TSS) within 1 kb of peak centres. The genomic regions enrichment annotations tool (GREAT) was used for gene ontology (GO) pathway analysis of significant differentially accessible regions. GREAT annotates proximal and distal chromatin regions to their target genes based on prior annotations from the literature (McLean *et al*. 2010). All differentially accessible peaks passing cut-offs were used as a background to calculate the statistical significance of enrichment using Fisher’s exact test. GO biological pathways with adjusted p-value < 0.05 were ranked by enrichment, and only terms that contained between 10 and 500 genes were retained. GO terms were also evaluated for redundancy, where redundancy was defined as 40% overlap of the genes included in the GO term. Finally, the most highly enriched, non-redundant terms were plotted by significance. Significant DAR from each genotype were sub-grouped by log_2_FC > 1 or log_2_FC < −1. Each subset was analyzed with GREAT as previously described, with the exception that all significant GO terms are displayed for differentially accessible regions where chromatin accessibility decreased in PPM1A^+^ cells. All significantly differentially accessible regions in each genotype were analyzed using QIAGEN Ingenuity Pathway Analysis (IPA) software (Krämer *et al*. 2014). IPA used Fisher’s exact test with the Benjamini-Hochberg correction, and only terms with FDR <0.05 were retained. Terms classified as ‘Disease’, ‘Function’ or ‘Upstream Regulator’ had absolute z-score > 2.

### Transcription factor binding motif analysis

Transcription factor binding motif (TFBM) analysis was conducted on all significant differential ATAC-seq peaks in WT, ΔPPM1A and PPM1A^+^ cells upon Mtb infection. Transcription factor motif enrichment within the sequences was analyzed using the R/Bioconductor package *universalmotif* (Tremblay and Nystrom 2021) and the JASPAR 2022 clustered set of non-redundant vertebrate motifs (Castro-Mondragon *et al*. 2017, 2022). For all sets of sequences tested for enrichment, the background set was a shuffled version of the foreground using k-let of 2. Motif hits on sequences were recorded if they generated a p-value less than 0.0001. Enrichment was calculated with the formula:

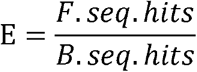

where E is the calculated enrichment, F.seq.hits represents the number of foreground sequences with at least one motif hit, and B.seq.hits represents the number of shuffled background sequences with at least one motif hit. This simple formula is appropriate considering that the number and length of sequences in the background and the foreground are equal. Enrichment significance was assessed by Fisher’s exact test and p values were corrected to generate q values using the False discovery rate method. Only motifs with E of at least 2.5 and q value less than 0.001 were considered enriched. For motif clusters, the most similar motif from JASPAR was identified using the compare_motifs function of *universalmotif*.

### RNA-seq library preparation

Macrophages were harvested as described for ATAC-seq. A third biological replicate was included for RNA-seq. mRNA extraction was performed using the Aurum™ Total RNA Mini Kit (Bio-Rad) as per the manufacturer’s protocols. All samples were incubated in lysis buffer for 20 minutes to inactivate bacteria in Mtb-infected samples. After extraction, the quality of mRNA was assessed using the Agilent RNA 6000 Nano LabChip Bioanalyzer. cDNA libraries were generated from mRNA samples with the NEB Ultra II Directional polyA mRNA library prep kit for Illumina. cDNA libraries were then sequenced with the Illumina NovaSeq platform for a total of 100 bp paired-end reads per mRNA sample to reach a target of 30-50 million total reads per sample. cDNA library preparation and sequencing were performed by The Centre for Applied Genomics, The Hospital for Sick Children, Toronto, Canada. Fasta sequence files were analyzed for quality using fastQC v0.11.9 (*FastQC* 2015). Remaining adaptor sequences and low-quality sequence fragments were removed using fastp v0.20.1 with the parameters --length_required 25, --cut_tail, cut_tail_window_size 4, --cut_tail_mean_quality 20, --disable_quality_filtering, -- overrepresentation_analysis. (Chen *et al*. 2018). Filtered reads were aligned to the human hg37 genome using STAR v2.7.5a, with the parameters --outReadsUnmapped Fastx, -- outFilterMatchNminOverLread 0.50, --outFilterScoreMinOverLread 0.50 --outSAMmapqUnique 40, -- outFilterMultimapNmax 1 (Dobin *et al*. 2013) and duplicate reads were marked with Picard (Picard Tools). Subread featureCounts was used to assign reads to gene exons and to summarize counts at the gene level (Liao, Smyth and Shi 2014). Represented genes were annotated using biomaRt (Smedley *et al*. 2009).

### Normalization and differential gene expression analysis

Differential expression analysis was performed using edgeR with glmQLFTest function for each comparison as described for ATAC-seq (Robinson, McCarthy and Smyth 2010). To be retained in the analysis, each gene had counts greater than 10 in at least 2 samples. Qualifying genes were analyzed with TMM normalization and batch effects were removed with RUVseq using RUVr k=3 (Risso *et al*. 2014; Peixoto *et al*. 2015). Genes with log_2_CPM (counts per million) less than zero were not included for further analysis. Differentially accessible genes with Benjamini-Hochberg adjusted P value (B-H adj. p-value), or FDR, less than 0.05 and absolute log_2_FC value of at least 1 were considered significant. Heatmaps were generated using the complexHeatmap package and PCA plots were generated with plot PCA function (Gu, Eils and Schlesner 2016).

### RNA-seq Gene Set Enrichment Analysis

Gene set enrichment analysis (GSEA) was conducted with all differentially expressed genes, using the R clusterProfiler package with the parameters minGSSize = 10, maxGSSize = 6000, pvalueCutoff = 1.0, pAdjustMethod = “BH”and Eps = 0 (Wu *et al*. 2021). Only gene sets that were GO biological pathways were considered for further analysis as ATAC-seq pathway analysis was limited to GO biological pathways. GO biological pathways with FDR less than 0.05 that contained between 10 and 500 genes were ranked by normalized enrichment score (NES) and considered significant if the NES was greater than 1, or less than −1. The GO biological processes that the most significantly enriched in WT cells (‘Type I IFN response’, and ‘Defense Response to Virus’) were selected for further analysis. Gene set enrichment plots were used to determine the leading-edge genes enriched in each pathway. Heatmaps of the leading-edge genes were generated using normalized log_2_FC differential expression values were median centred across all samples of the dataset. All significantly differentially expressed genes in each genotype were also analyzed using QIAGEN IPA software, with the same parameters used for ATAC-seq analysis (Krämer *et al*. 2014).

### Integration of ATAC-seq and RNA-seq Data

Genes that were both differentially accessible and expressed were determined by matching the genes that CA peaks were annotated to, with DEG. For DEG with multiple CA peaks, the peak closest to the TSS of the gene was selected to be the match for further analysis. A maximum distance of 50 kb between peak center and the gene TSS was set. Matching genes were considered significant if they had FDR less than 0.05 and absolute log_2_FC > 1 for both CA and gene expression. A separate dataset including log_2_FC was generated for all significant overlapping DAR and DEG in WT, ΔPPM1A and PPM1A^+^ cells for comparison. For Figure 6C overlapping genes were ranked by log_2_FC gene expression, and the top 10 upregulated or downregulated genes in each genotype were selected. Log_2_FC values were displayed for all of these genes in each genotype, even if they were not significant.

## ACKNOWLEDGMENTS

We thank members of the Sun lab for insightful discussions and the Common Equipment and Technical Service (CETS) at the University of Ottawa for expert technical assistance. We would also like to thank Karen Ho and Sergio Pereira at The Centre for Applied Genomics, The Hospital for Sick Children, Toronto, Canada for assistance with next-generation sequencing, and Compute Canada for access to the Cedar high-performance computing cluster. This work was supported by grants from the Canadian Institutes of Health Research (CIHR) PJT-162424 and the National Sanitarium Association Scholar’s Program to J.S, a CIHR operating grant (MOP 119458) to A. B., and the University of Ottawa Faculty of Medicine Translational Research Grant to J.S. and G.G.A. K.M. and R.E.H were supported by graduate scholarships from the University of Ottawa Centre for Infection, Immunity and Inflammation.

## DATA AVAILABILITY

All ATAC-seq and RNA-seq data will be made publicly available on the NCBI Gene Expression Omnibus (GEO) and an GEO accession number will be provided in a revised version of this preprint.

## CONTRIBUTIONS

Conceptualization, methodology, and visualization, K.M., R.E.H., S.B., G.G.A., A.B., and J.S.; investigation, K.M., R.E.H., S.B., and A.B.; writing – original draft, K.M. and J.S.; writing – review & editing, K.M., R.E.H., S.B., G.G.A., A.B., and J.S.; supervision, funding acquisition, and resources: G.G.A, A.B. and J.S.

## SUPPLEMENTARY DATA (FIGURES S1-9)

Figure S1. Optimal Mtb infection conditions in THP-1 macrophages.

Figure S2. ATAC-seq quality control metrics.

Figure S3. GREAT gene association graphs.

Figure S4. Ingenuity Pathway Analysis (IPA) of differentially accessible regions (DAR).

Figure S5. Quantification of enriched Transcription Factor Binding Motifs (TFBM).

Figure S6. Principal Component Analysis (PCA) of all expressed genes.

Figure S7. Gene Set Enrichment Analysis (GSEA) plots.

Figure S8. Ingenuity Pathway Analysis (IPA) of differentially expressed genes (DEG).

Figure S9. Quantification and characterization of concordant DAR and DEG.

