## Supplemental Data (Figure S1-9) for "Transcriptome and chromatin accessibility mapping reveals a type I Interferon response triggered by *Mycobacterium tuberculosis* infection"

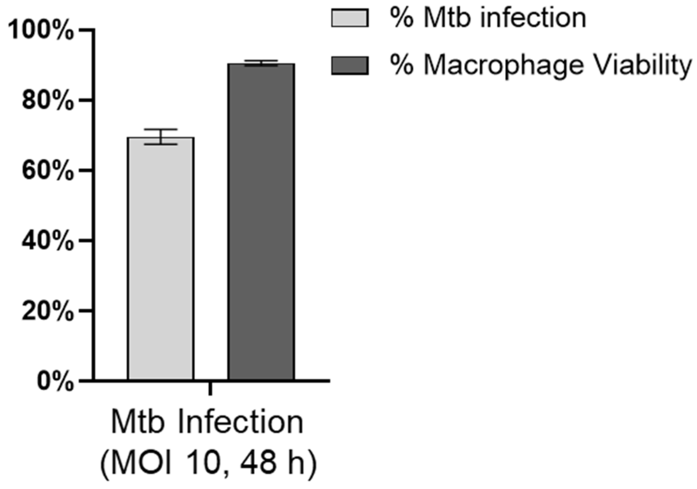


**Figure S1. Optimal Mtb infection conditions in THP-1 macrophages.** THP-1 macrophages were infected with Mtb mc$^2$6206 expressing green fluorescent protein (GFP) at an MOI of 10. Extracellular non-phagocytosed bacteria were removed by washes at 16 h post-infection and infection was continued until 48 h post-infection. Then, macrophage viability was assessed using the BD Horizon fixable viability stain 780 and analyzed by flow cytometry. The percent of macrophages infected with Mtb was determined by analyzing the GFP signal using flow cytometry.

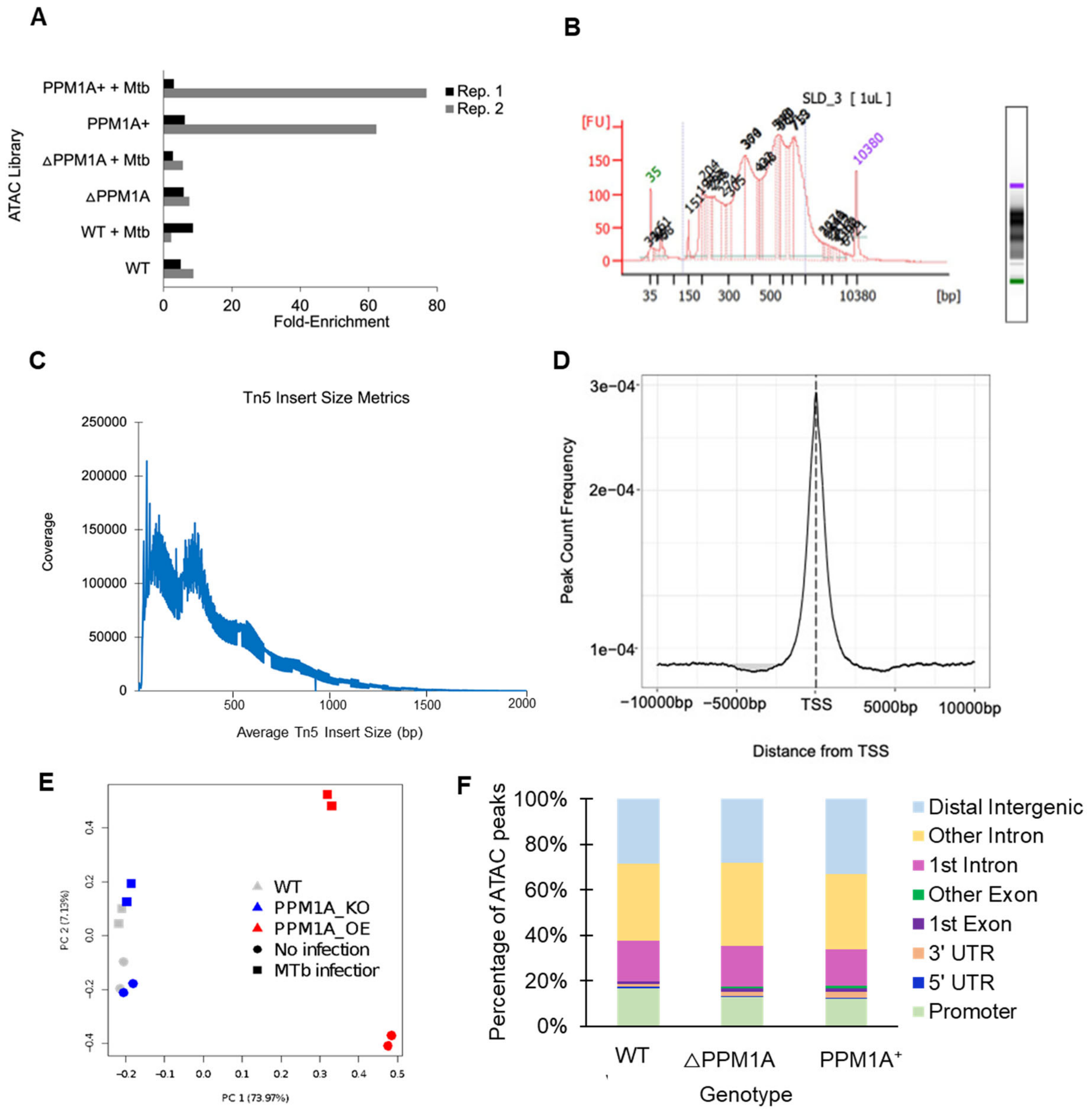


**Figure S2. ATAC-seq quality control metrics.** **(A)** DNA in chromatin regions known to be either accessible or inaccessible in THP-1 cells was measured in non-infected and Mtb-infected WT, △PPM1A and PPM1A$^{+}$ macrophages was measured by qPCR using gene specific primers. **(B)** Bioanalyzer electropherogram showing a representative sample where the average fragment size was 478 bp, and the concentration was 5.1 ng/µl. **(C)** PicardTools InsertSizeMetrics was used to measure the number of Tn5 transposed fragments compared to their length in base pairs. **(D)** Average profile of all ATAC-seq peaks. The genomic distance in base pairs of each peak from its annotated gene TSS was plotted against the number of reads of each peak using ChIPseeker. **(E)** Principal component analysis (PCA) of all ATAC-seq peaks. Read coverage over all peaks demonstrated that genotype and infection status were the largest and second largest contributor to variance in the dataset, respectively. Peaks from biological replicates also clustered together, confirming their similarity. **(F)** Genomic features of DAR. Distribution of different genomic features of DAR were determined using ChIPseeker in WT, △PPM1A and PPM1A$^{+}$ cells upon Mtb infection. The majority of DAR in all genotypes were found in "Other introns" and "Distal intergenic" regions.

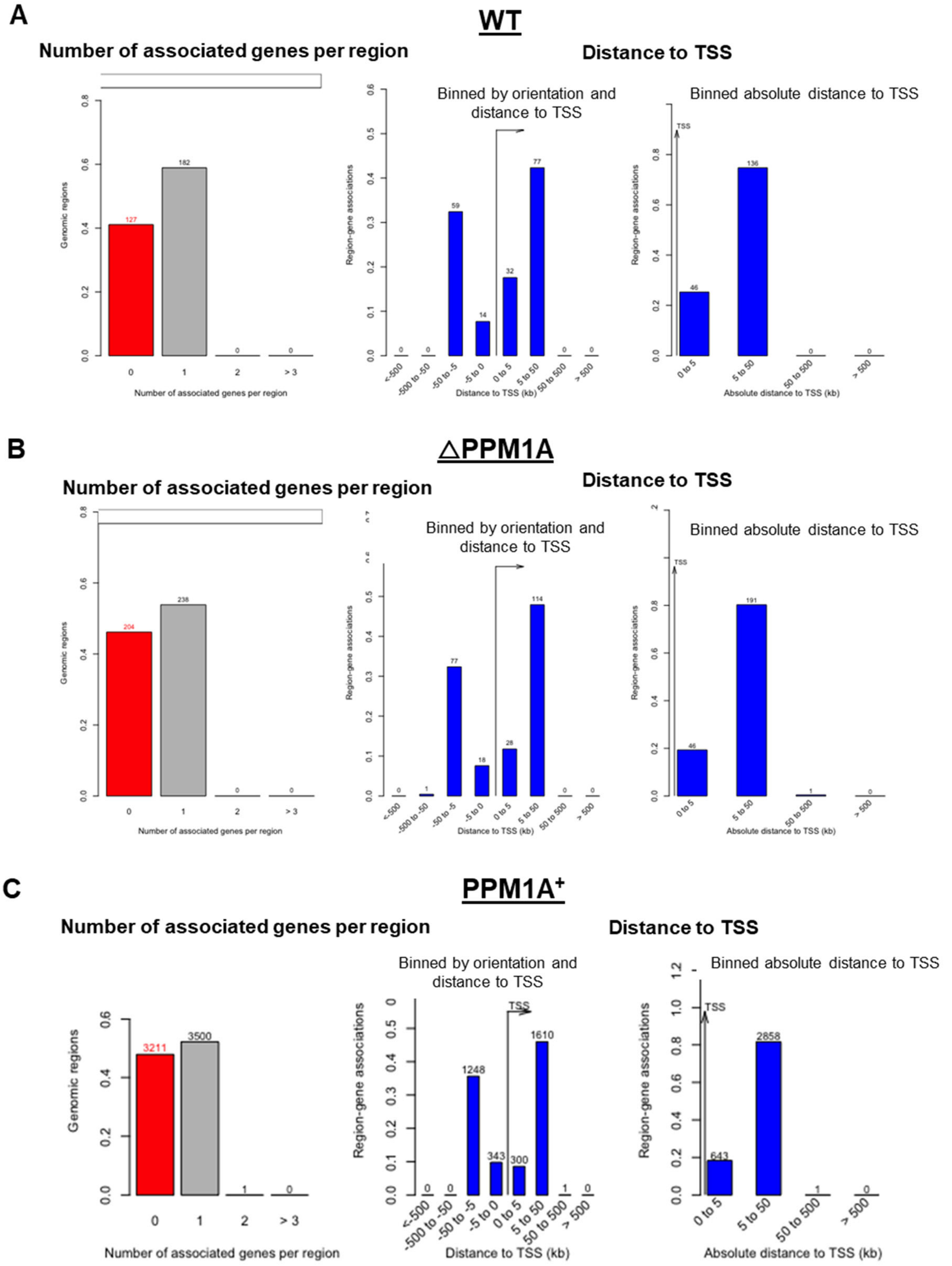


**Figure S3. GREAT gene association graphs. (A-C)** The number of associated genes in each DAR were determined by the gene association rules defined by GREAT (left). The majority of DAR in all genotypes were associated with one gene. Distance to TSS graphs are first displayed where TSS were categorized by whether they were upstream or downstream of TSS (middle), and then displayed where only the absolute genomic distance is considered (right).

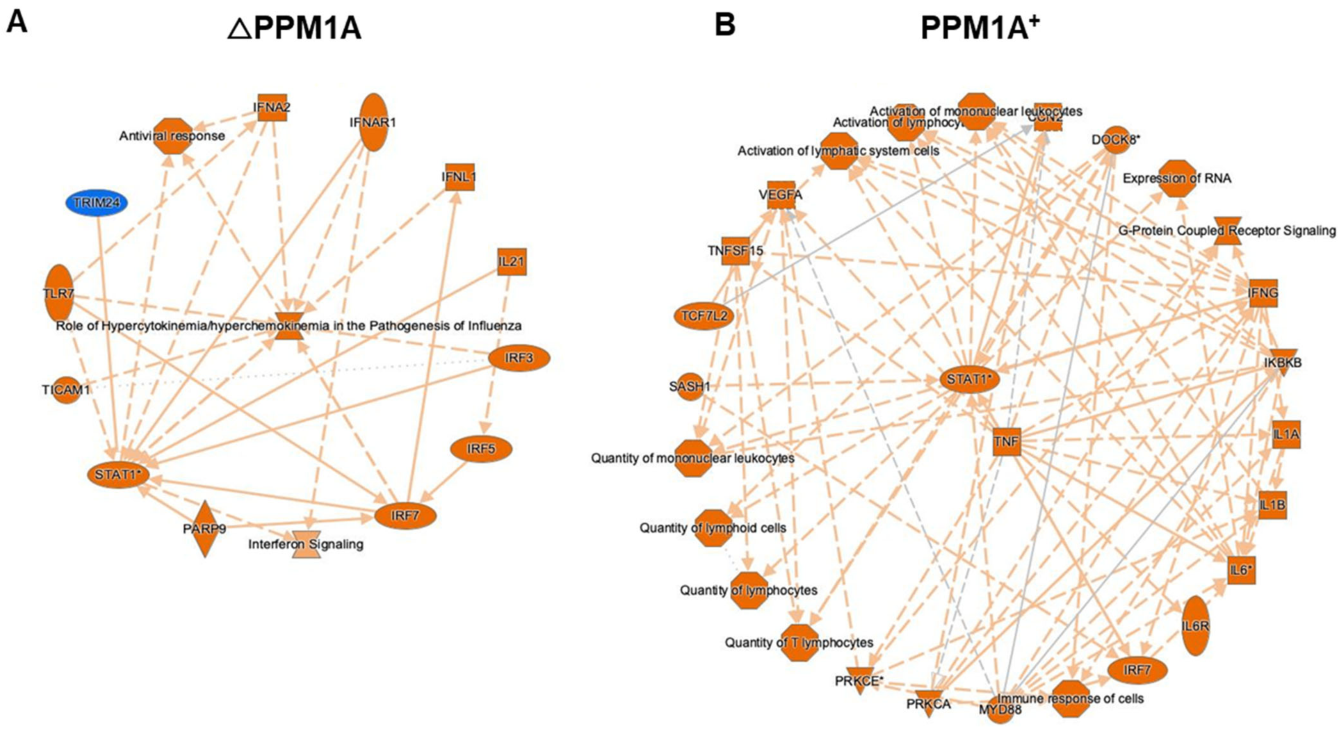


**Figure S4. Ingenuity Pathway Analysis (IPA) of differentially accessible regions (DAR). (A, B)** IPA was conducted on all significant DAR for **(A)** △PPM1A and **(B)** PPM1A$^+$ macrophages following Mtb infection, and the radial summary graph is shown.

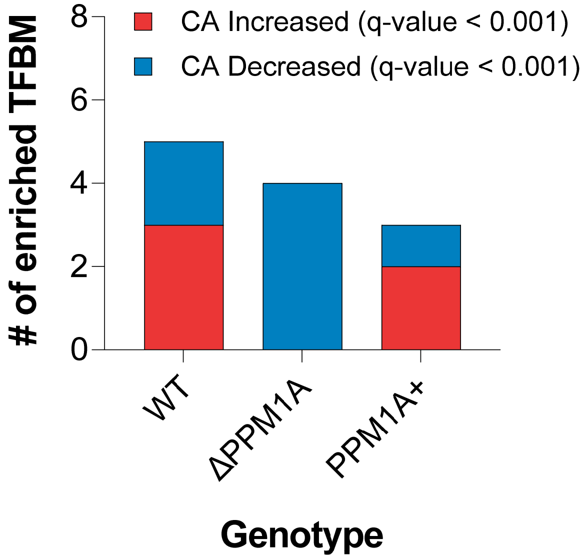


**Figure S5. Quantification of enriched Transcription Factor Binding Motifs (TFBM).** Total numbers of enriched TFBM found in DAR in WT, △PPM1A and PPM1A$^{+}$ macrophages following Mtb infection.

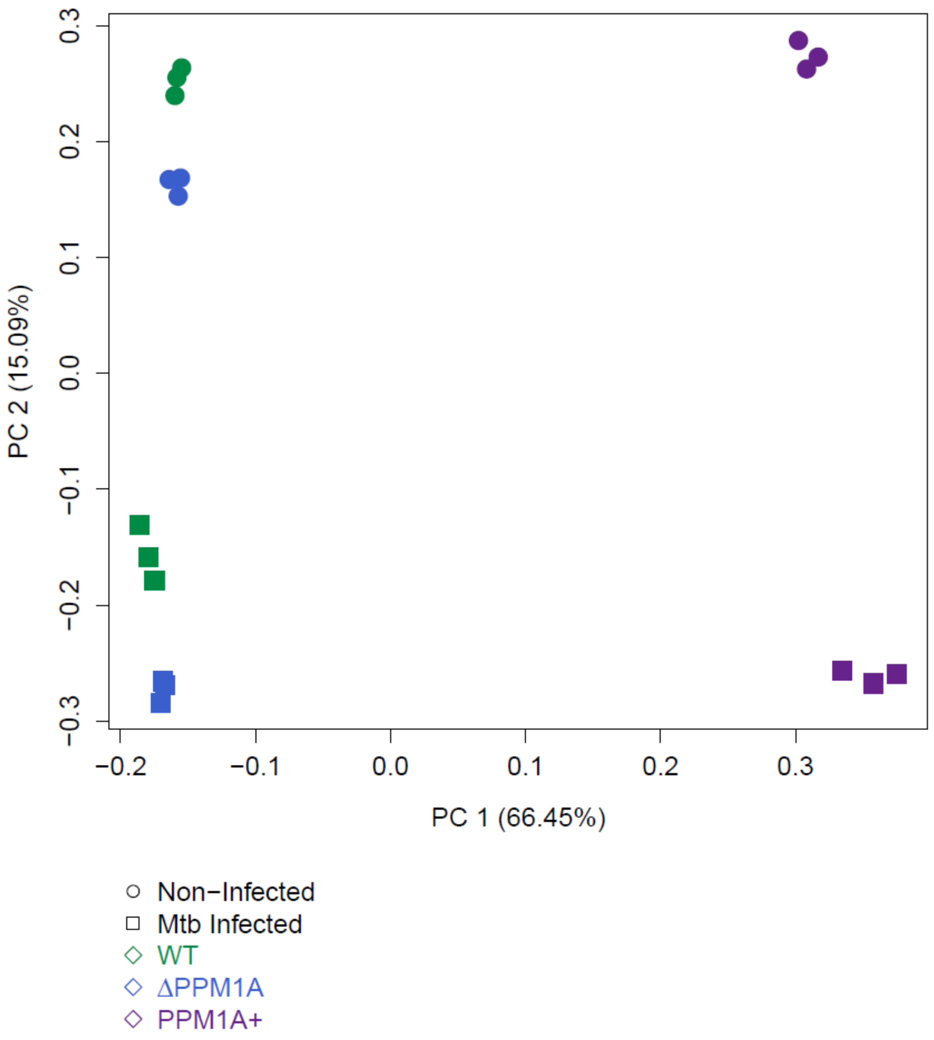


**Figure S6. Principal Component Analysis (PCA) of all expressed genes.** PCA analysis of total gene expression data displays that genes separate and cluster together based on the first and second principal components which are genotype and Mtb infection status, respectively. Genotype accounts for 66% of the variance observed in all expressed genes, and Mtb infection accounts for 15%.

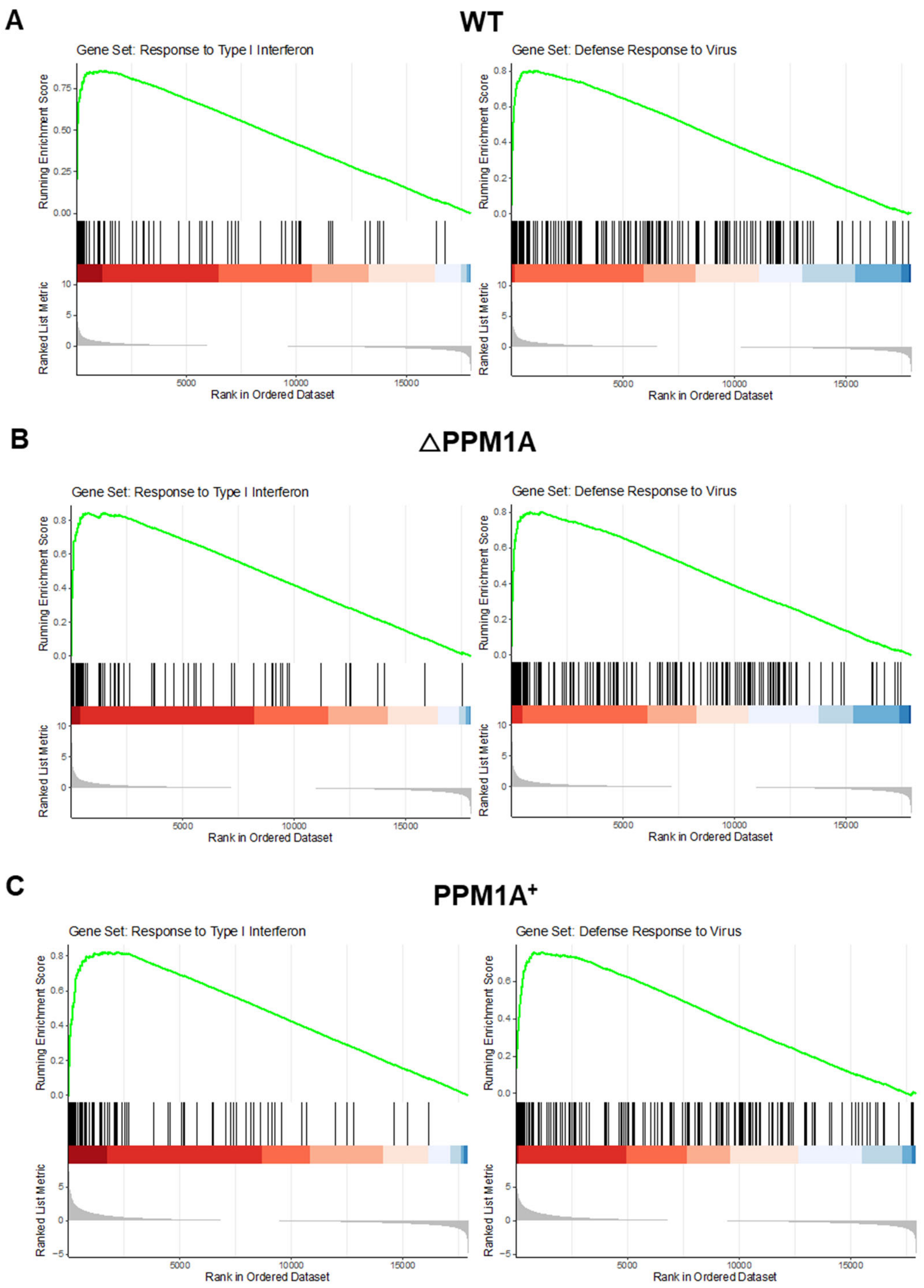


**Figure S7. Gene Set Enrichment Analysis (GSEA) plots. (A-C)** GSEA was performed in R using the clusterProfiler package against MsigDB 7.5.1 Hallmark, C2, and C5 gene sets. The p-value was adjusted using the Benjamini-Hochberg procedure. Genes with low expression in all samples were removed from the analysis. GSEA was conducted on Mtb-infected and non-infected WT, △PPM1A, and PPM1A$^{+}$ macrophages. The total height of the curve corresponds to the overall enrichment score (ES), with the leading-edge genes appearing prior to the peak score.

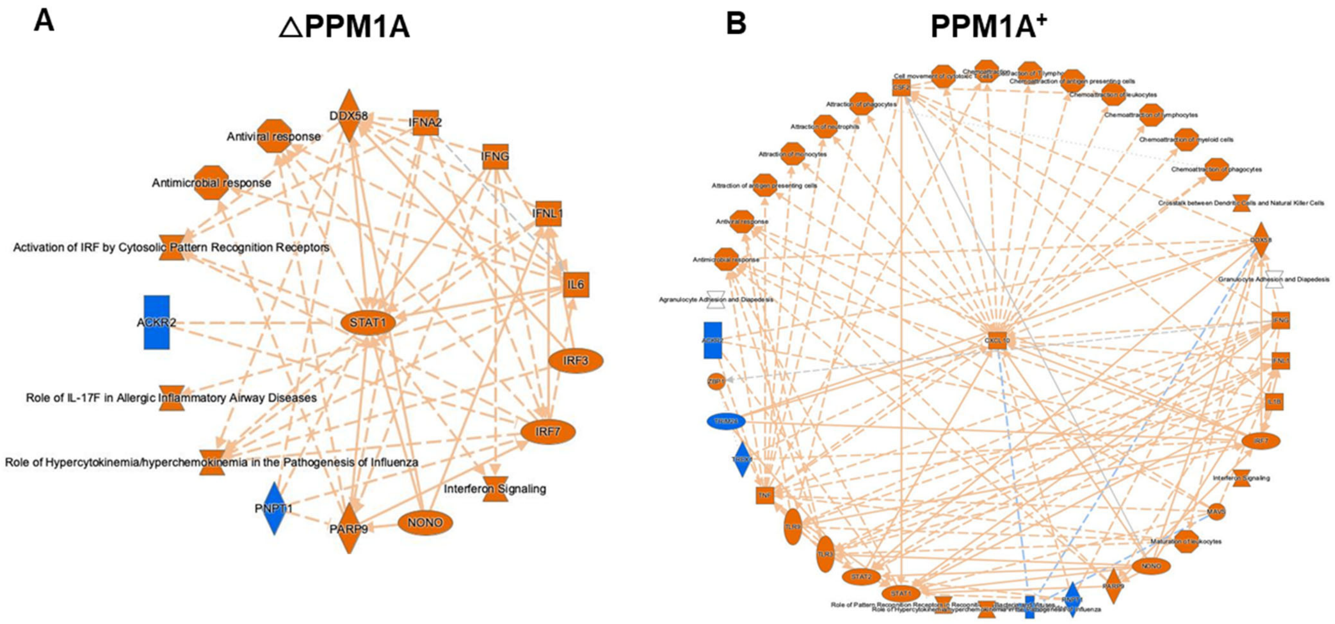


**Figure S8. Ingenuity Pathway Analysis (IPA) of differentially expressed genes (DEG). (A, B)** IPA was conducted on all significant DEG for **(A)** △PPM1A and **(B)** PPM1A$^+$ macrophages following Mtb infection and the radial summary graphs are shown.

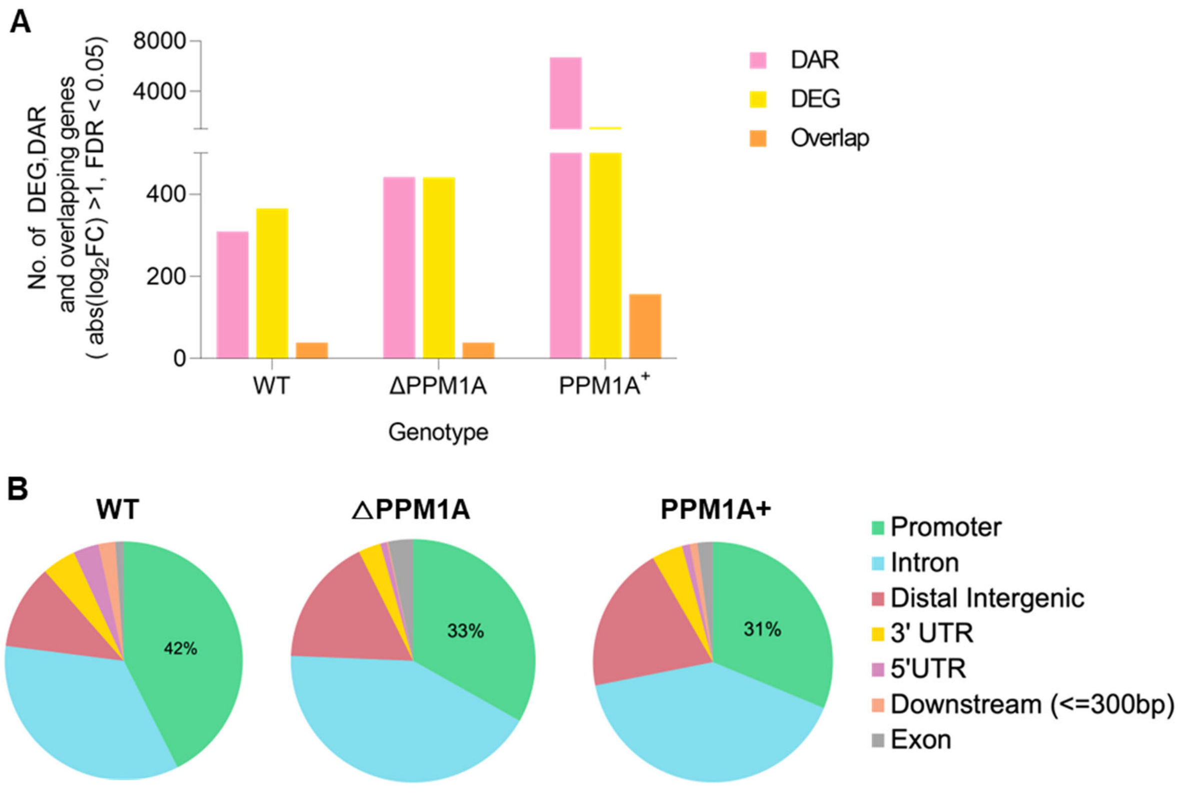


**Figure S9. Quantification and characterization of concordant DAR and DEG. (A)** Total numbers of all significant DAR (pink), DEG (yellow) and concordant genes (orange) were quantified for comparison. **(B)** Genomic features of DAR with concordant DEG. DAR with concordant DEG in WT, △PPM1A and PPM1A$^{+}$ macrophages were annotated to genomic features using ChIPseeker. The majority of DAR in all genotypes were found in "Introns" and "Distal intergenic" regions.
